# Radiation resistant cancer cells enhance the survival and resistance of sensitive cells in prostate spheroids

**DOI:** 10.1101/564724

**Authors:** Pavitra Kannan, Marcin Paczkowski, Ana Miar, Joshua Owen, Warren W. Kretzschmar, Serena Lucotti, Jakob Kaeppler, Jianzhou Chen, Bostjan Markelc, Leoni A. Kunz-Schughart, Adrian L. Harris, Mike Partridge, Helen Byrne

## Abstract

Intratumoural heterogeneity contributes to local tumour recurrence and variable responses to radiotherapy in prostate cancer. Despite the multiclonal nature of the disease, tumour control probability for conventional treatment plans is modelled on the assumption that tumour cells in the target region respond identically and independently. Here, using tumour cell subpopulations with different radiation sensitivities from prostate tumour cell lines, we show that radiation resistant cells enhance the survival and radiation resistance of radio-sensitive cells in spheroids but not in monolayer culture. Mathematical modelling indicates that these phenotypic changes result from both competitive and antagonistic cellular interactions in spheroids. Interactions mediated by oxygen constraints define the spatial localisation of the cell populations in spheroids and in xenografts, while those mediated by paracrine signals further modify the microenvironment. Our results show new mechanisms of radiotherapy resistance mediated by cellular interactions and by the microenvironment.

## INTRODUCTION

External beam radiotherapy, combined with androgen deprivation, is commonly used to treat patients with localised prostate cancer^1^. Despite advances in radiotherapy that have led to more targeted therapies with reduced toxicity, local recurrence remains a challenge to manage in treatment because patients with similar clinical risk parameters (such as Gleason score, PSA levels, and staging) have variable outcomes and mortality rates^2^. One factor that may contribute to variable responses and recurrence is intratumoural heterogeneity^3^. Primary and metastatic prostate lesions have been found to contain multiple genetically and clonally distinct foci, with more than 80% of lesions containing at least 1 disease focus^3–6^. Clonal dynamics can change during therapy and may lead to the maintenance of intratumoural heterogeneity after therapy, facilitating recurrence^7–9^.

Interactions between cell populations can also influence tumour growth and therapeutic outcome^10–12^. Similar to interactions between species in other ecological systems^13^, interactions in tumours, in which one cell population influences another, can be manifested indirectly or directly. Indirect interactions occur through microenvironmental pressures, such as hypoxia, which force cells to adapt and/or compete for resources^14–17^. Direct interactions occur through communication between cell populations (i.e., paracrine signalling). Secreted proteins between tumour-stromal cells^18–20^, or between tumour-tumour cells^14,21–27^, have been shown to increase growth, metastasis, and resistance to chemotherapy in melanoma, breast, prostate, and small cell lung cancers. However, little is known about whether interactions (indirect and/or direct) between tumour cell clones affect radiotherapy response.

Conventional treatment plans for external beam radiotherapy, however, do not directly account for intratumoural interactions. The mathematical model used to predict tumour control probability assumes that all tumour cells in the target region have the same radiation sensitivity. Furthermore, cell kill events are modelled as a Poisson process, with subsequent events occurring independently of one another. Simulations of tumour response to radiotherapy indicated that modulating the radiation dose to account for intratumoural heterogeneity improves local control and reduces recurrence in silico^28,29^. However, these models do not account for interactions (either indirect or direct) between populations of radio-sensitive and radio-resistant cells, and whether these interactions impact radiotherapy response. Furthermore, experimental evidence demonstrating the impact of cellular interactions on radiation response is lacking.

We therefore sought to determine whether indirect and/or direct interactions occur between tumour cell populations with different radiation sensitivities, and whether these interactions affect bulk tumour growth kinetics and radiation response. Control and radiation resistant cell populations from prostate cell lines were combined as homogeneous (control or resistant alone) and mixed (control and resistant together) populations. Phenotypic changes in growth before and after radiation were measured using 2D and 3D biological assays. Mathematical modelling was used to determine whether changes in growth before and after treatment could result from cellular interactions and to determine the nature (e.g., competition vs cooperation) of these interactions.

## RESULTS

### Inclusion of radiation resistant cells increases growth kinetics in 3D but not in 2D

Control and radiation resistant cell populations from two prostate cell lines, PC3^30^ and DU145^31^, were used as a model to investigate whether cellular interactions between mixed populations could alter bulk tumour growth. After confirming that control and resistant populations had inherently different radiation sensitivities in both cell lines (SI Fig 1) we measured the growth of homogeneous and mixed (seeded 1:1 ctrl:res) populations as 2D monolayers and 3D spheroids. Inclusion of 50% resistant cells did not detectably change the growth rates of PC3 or DU145 monolayers (Fig 1a). In contrast, the volumes of mixed PC3 spheroids increased significantly by > 30% beginning at day 10 (P < 0.001) and that of mixed DU145 spheroids by > 22% beginning at day 15 (P < 0.001, Fig 1b). Because the change in growth could result from the resistant population outgrowing the control one, we generated cell populations stably expressing GFP (control) or DsRed (resistant), grew mixed spheroids (seeded 1:1 ctrl:res), and dissociated them to quantify the proportions of each cell population over time using flow cytometry (SI Fig 2). On day 15 of growth, resistant cells isolated from mixed PC3 spheroids comprised 60.9 ± 9.6% of total cells, while those isolated from mixed DU145 spheroids comprised 22.7 ± 12.2% of total cells (Fig 1c). These results show that inclusion of resistant cell lines increases the growth of mixed tumour populations in 3D but not in 2D, and that the enhanced population may be control cells.

**Figure 1.**
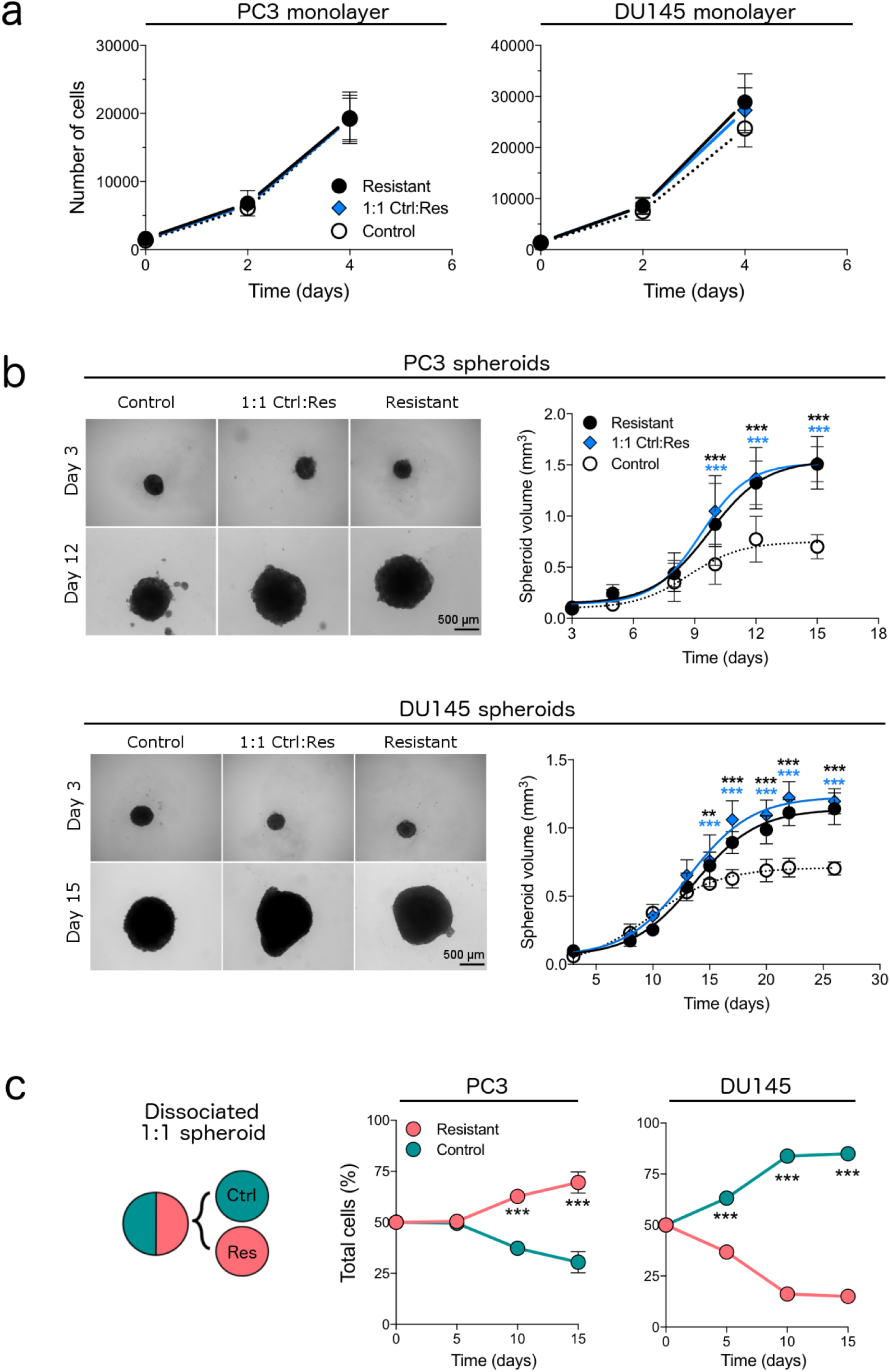
Inclusion of radiation resistant cells increases bulk growth of mixed tumour cell populations in 3D but not 2D for two prostate cancer cell lines. **(a)** Growth curves of PC3 and DU145 monolayer cells comprising radiation sensitive (control), radiation resistant (res), or 1:1 mixtures of control:resistant (ctrl:res). **(b)** Growth curves and representative images of 3D spheroids comprising control, resistant, or 1:1 mix of ctrl:res cells. **(c)** Proportions of control and resistant populations measured over time using flow cytometry from spheroids seeded as 1:1 mixture. Data represent mean ± SD from 3 independent experiments except for **b**, for which data represent mean ± SD of 12 spheroids pooled from 2 independent experiments. *** *P* < 0.001, as determined by 2-way ANOVA followed by Bonferroni correction for multiple testing.

### Mathematical modelling predicts that enhanced growth kinetics result from cellular interactions between mixed populations

To differentiate between changes that result from a simple difference in proliferative capacity versus those that result from cellular interactions, we developed and applied a mathematical model using Lotka-Volterra-type interactions. This model extends the logistic growth models used to describe the growth of the homogeneous spheroids by introducing cellcell interaction terms that describe interactions found in evolutionary biology: mutualism (mutually beneficial or cooperation), antagonism (one population benefits while the other is negatively affected) and/or competition (mutually detrimental).^13^ As a first step, the logistic growth model was fitted to growth curves of homogeneous spheroids to estimate three parameters for each tumour cell population: the initial growth rate r, the carrying capacity *K* (or maximum spheroid volume at steady state), and the initial volume *V_0_* (Fig 2a). The parameter values *r* and *K* were higher for the resistant population than for the control population in PC3 spheroids. In DU145 spheroids, the growth rate *r* of the resistant cells was lower than that of the control cells while the carrying capacity *K* was higher (Fig 2b); values of *V_0_* were not different between cell lines (SI Table 1). These parameter values were subsequently held fixed. The interaction parameters, *λ_C_* and *λ_R_*, were then estimated by fitting the Lotka-Volterra model to data from the growth curves of the mixed spheroids (Fig 2c). The nature of interactions between the two cell populations depends on the sign of each of the interaction parameters *λ_C_* and *λ_R_*, resulting in three main types of interactions: competition (*λ_C_* < 0 and *λ_R_* < 0), antagonism (*λ_C_* < 0 < *λ_R_* or *λ_C_* > 0 > *λ_R_*), or mutualism (*λ_C_* > 0 and *λ_R_* > 0).

**Figure 2.**
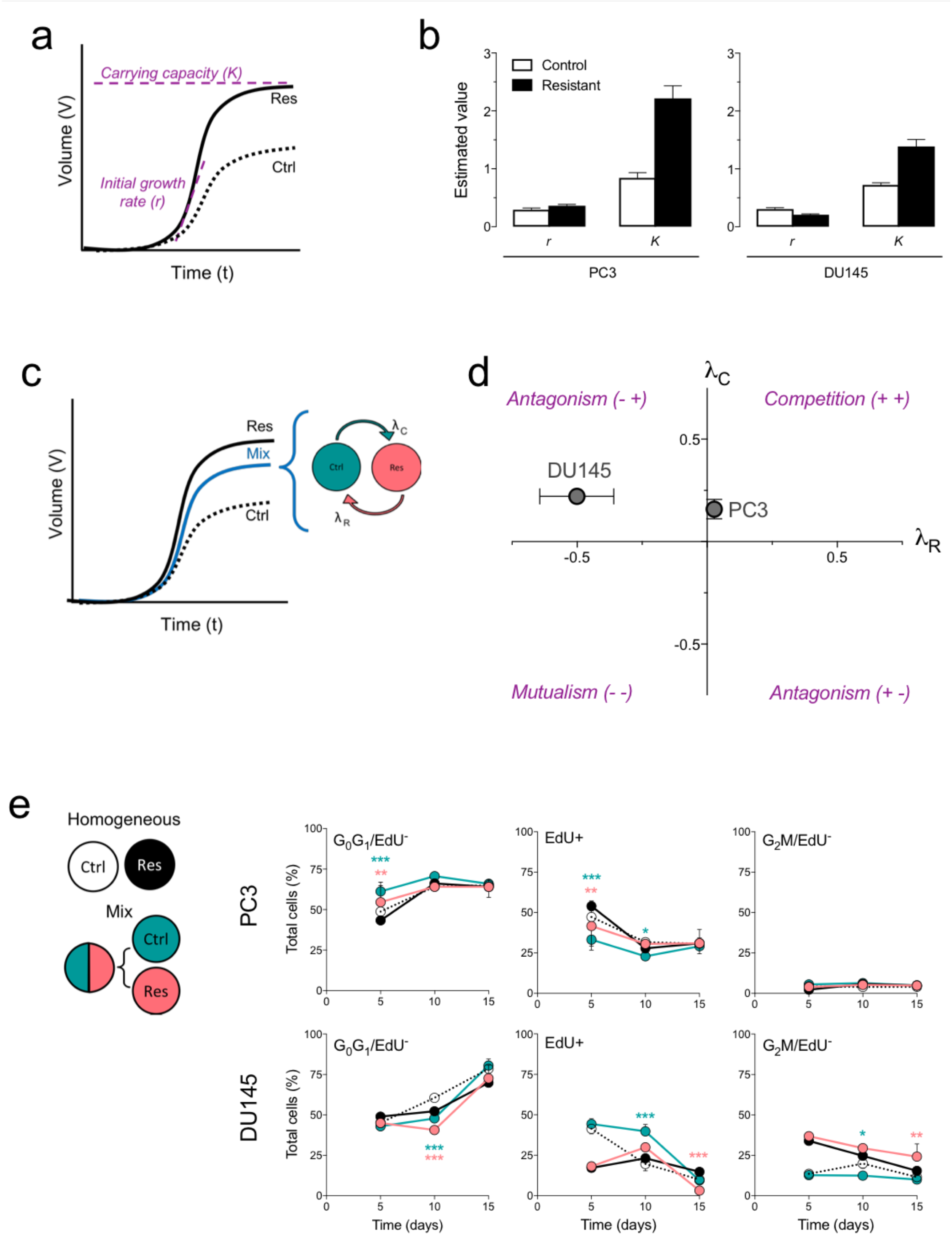
Mathematical modelling predicts the nature of cellular interactions in mixed prostate spheroids. **(a)** Logistic model of spheroid growth, with growth rate *r* and carrying capacity *K* (the limiting volume of the spheroid) of tumour cell populations. **(b)** Estimates of growth parameters for control and resistant cells calculated by fitting the logistic model to growth curves of homogeneous PC3 and DU145 spheroids. Error bars represent confidence intervals. **(c)** Schematic of Lotka-Volterra model to measure growth curve of mixed spheroids, in which tumour cells interact with one another. The parameter *λ_C_* indicates the effect of controls on resistant, while the parameter *λ_R_* indicates the effect of resistant on control. **(d)** Estimated interaction parameters using biological data of PC3 and DU145 spheroids containing 1:1 mixture of ctrl: res cells. The nature of interaction, i.e., competition vs antagonism, depends on the sign of the interaction parameters. Error bars represent confidence intervals. **(e)** Experimental determination of cell cycle distribution for control and resistant cell populations isolated from mixed and homogeneous spheroids. Data represent mean ± SD from 4 independent experiments. * *P* < 0.05, ** *P* < 0.01, and *** *P* < 0.001, as determined by 2-way ANOVA followed by post-hoc t-tests with Bonferroni correction for multiple testing.

To reliably infer the interaction parameters in our experimental data, we calculated the uncertainty associated with inference using synthetic growth curves of mixed spheroids with known interaction parameters (*λ_C_* = 0.5 and *λ_R_* = 0.5) and three levels of experimental noise (5, 10, and 20%). Interaction parameters estimated by fitting volume measurements to the Lotka-Volterra model resulted in large uncertainties (SI Fig 3a), whereas interaction parameters estimated using both volume measurements and the ratio of the mixed populations substantially decreased uncertainty (SI Fig 3b). Using both volume measurements and cellular proportions from our experimental data (Fig 1), we deduced competitive interactions in PC3 spheroids and antagonistic interactions in DU145 spheroids (Fig 2d).

### Experimental data validate predictions from mathematical models of cellular interactions

To test whether the mathematical predictions could be supported by experimental data, we measured changes in cell cycle turnover and survival of each population isolated from homogeneous and mixed (1:1 ctrl:res) spheroids over time (Fig 2e). The uptake of the nucleoside analogue 5-ethynyl-2 deoxyuridine (EdU) was detected both in cells proliferating in S phase and in cells that previously took up EdU in S phase and then cycled into G0G1 during the incubation time; we thus gated cells based on presence/ absence of EdU staining to account for turnover (SI Fig 4). On day 5, control and resistant cells isolated from mixed PC3 spheroids with G0G1/ EdU^-^ staining increased by 21% and 22%, while EdU^+^ uptake reduced by 30% and 25%, respectively (Fig 2e). Despite this competitive effect on cell cycle, measurements of cell death (SI Fig 5a) for both populations in mixed spheroids were reduced at day 15, as determined by efluor-780 staining for live and dead cells (SI Fig 5b). These findings suggest that the two PC3 populations adversely affect each other’s growth during the early stages of spheroid growth, but eventually they reach an equilibrium that enhances their overall viability.

Cell cycle phases and survival were also altered in mixed DU145 spheroids. Control cells in mixed spheroids had a 10 % increase in EdU^+^ uptake at day 10 of growth (Fig 2e), and significant reductions in G0G1/ EdU^-^ and G2M/EdU^-^ staining. In contrast, resistant cells in mixed spheroids had a significant increase in G2M/EdU^-^ staining and an 80% reduction in EdU^+^ uptake on day 15 (Fig 2e), but a decrease in cell death (SI Fig 5c). These results suggest that cells in DU145 behave antagonistically: the control cells gain a growth advantage, while the resistant cells are adversely affected. Our biological data thus confirm the mathematical predictions and indicate that for the cases considered, interactions, irrespective of the type (i.e., competitive, antagonistic), increase bulk tumour growth.

### Resistant cells increase bulk resistance and survival of control cells after radiation

To determine whether mixed populations had altered radiotherapy response, we irradiated homogeneous and mixed spheroids at a range of doses and measured their regrowth. Compared to PC3 spheroids comprising 100% control cells [SCP_50_, 5.17 Gy; 95% CI, 5.01-5.32 Gy], the dose required to control (or prevent regrowth of) 50% spheroids increased by 2 Gy with the inclusion of 10% resistant cells [SCP_50_, 7.21 Gy; 95% CI, 7.06-7.36 Gy] and by another 2 Gy with the inclusion of 50% resistant cells [SCP_50_, 9.56 Gy; 95% CI, 9.54-9.60 Gy] in mixed spheroids. Spheroids comprising 100% resistant cells had the highest required dose for control [SCP_50_, 11.14 Gy; 95% CI, 11.14-11.15 Gy] (Fig 3a). Inclusion of resistant cells shortened the time of spheroid regrowth after radiation. After 6 Gy radiation, mixed PC3 spheroids (1:1 ctrl:res) grew back faster than control spheroids (median days to regrowth: mixed = 18, control = 23; *P* < 0.001). All mixed spheroids regrew, while 25% of control spheroids did not (Fig 3b). Analysis of the cell proportions by flow cytometry revealed that 1: 1 mixed PC3 spheroids comprised 61 ± 5.6% and 79 ± 5.6% resistant cells on days 10 and 15, respectively (Fig 3c), suggesting that regrowth is driven by resistant cells. While survival of control cells isolated from mixed spheroids on day 15 was 80% higher than that of control cells isolated from homogeneous spheroids (SI Fig 6b), proliferation (of both current and cycled cells) dropped by 40% with an increase within G2M/EdU^-^, suggesting cell cycle arrest (SI Fig 7a).

**Figure 3.**
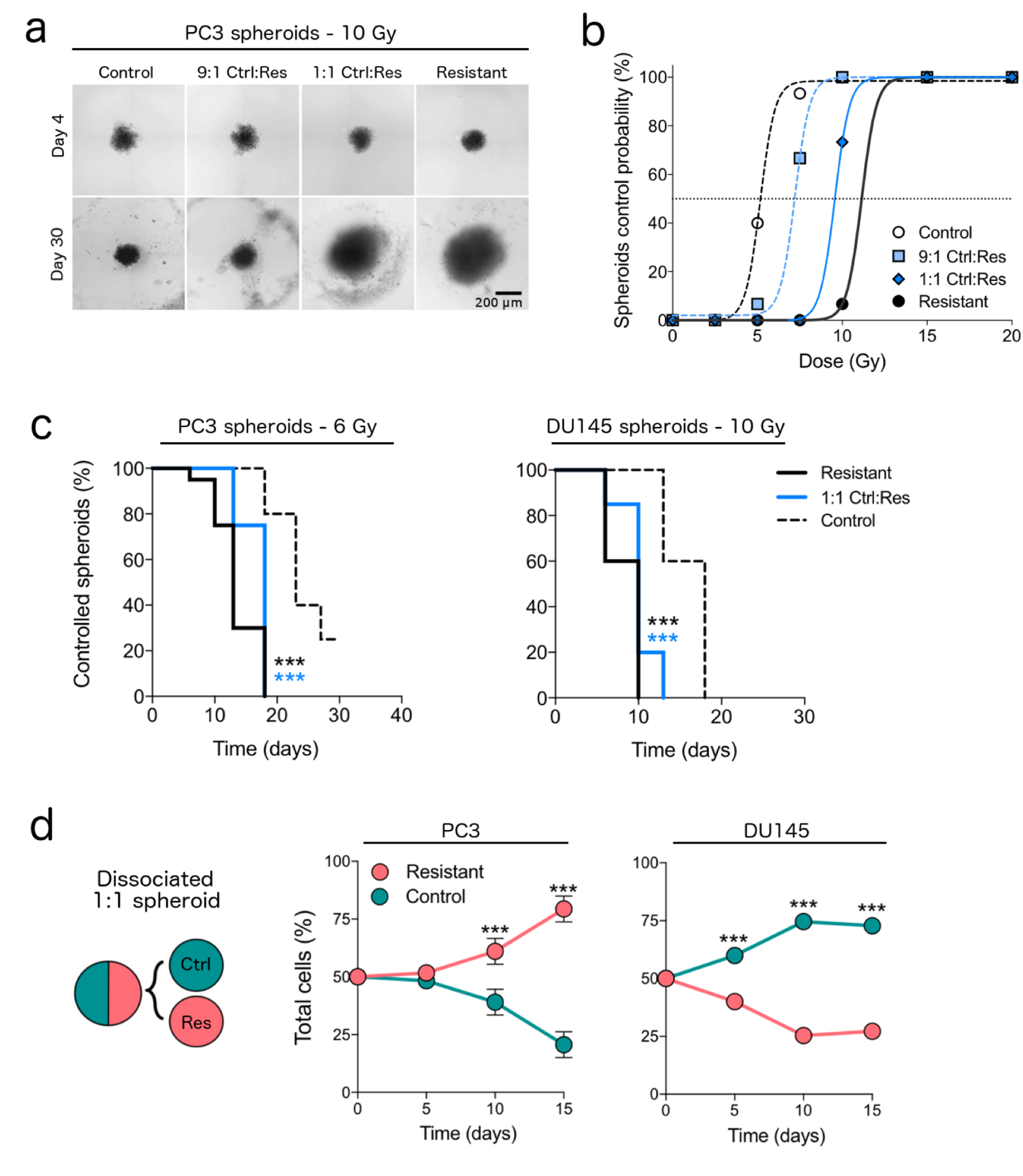
Radiation resistant cells increase bulk radiation resistance of mixed spheroids. **(a)** Images of irradiated PC3 spheroids seeded with 100% control cells, 9:1 ctrl:res, 1:1 ctrl:res or 100% resistant cells. **(b)** Control probability of irradiated spheroids comprising control, 9:1 ctrl:res, 1:1 ctrl:res, or resistant cells. Dashed line shows the 50% control probability that prevents regrowth. Data represent the percent of spheroids that grew back (n = 15 spheroids/group/radiation dose) from one biological experiment **(c)** Survival curves of irradiated spheroids seeded as control, 1:1 ctrl:res, or resistant. The experiment was performed on a smaller scale to replicate results of b. Data represent n = 20 spheroids/group. *** *P* < 0.001, as determined by Mantel-Cox survival analysis. **(d)** Proportions of control and resistant populations measured over time using flow cytometry from spheroids seeded as 1:1 mixture and irradiated at 6 or 10 Gy, respectively. Data represent mean ± SD from 4 independent experiments. *** *P* < 0.001, as determined by 2-way ANOVA followed by post-hoc t-tests with Bonferroni correction for multiple testing.

The regrowth phenotype was not specific to PC3 spheroids; a similar pattern was observed in DU145 spheroids treated with 10 Gy radiation (median days to regrowth: 1:1 mixed = 10, control = 18; *P* < 0.001; Fig 3c). This effect was not due to resistant cells dominating, as control cells comprised > 60% of the mixed spheroids (Fig 3d), and had a > 60% increase in survival (SI Fig 6c) in mixed spheroids, but an increase within G2M/EdU^-^ on day 10. Similarly, resistant cells in mixed spheroids had a 2-fold increase in survival (SI Fig 6c) and a 13% decrease within G0G1/ EdU^-^, but an increase within G2M/EdU^-^ on day 10 (SI Fig 7b). These results indicate that resistant cells can enhance the survival of the control population post radiation, and also that the control cells could contribute to post-radiation cell survival.

### The type of cellular interaction dictates regrowth time after radiation

We then investigated whether the type of interaction that occurs during growth could also alter the time of regrowth after radiotherapy. Using the Lotka-Volterra model, we simulated regrowth of mixed spheroids (seeded 1:1 in silico) after exposure to 6 Gy for a range of values of the interaction parameters *λ_C_* and *λ_R_*. Cell kill due to radiotherapy was modelled using the linear quadratic model^32^, with α and β values derived for each cell population (PC3 control and resistant) from clonogenic survival curves (SI Fig 1a); the oxygen dependence of *α* and *β* was not considered as the Lotka-Volterra model is spatially-averaged and oxygen levels are not included in the model. As expected, cell populations that compete with each other (*λ_C_* < 0 and *λ_R_* < 0) have the slowest regrowth time (50-200 days), while those that have mutualistic interactions (*λ_C_* > 0 and *λ_R_* > 0) have the fastest regrowth time (14-32 days) (Fig 4). Interestingly, within a specific interaction type, the numerical values of the interaction parameters *λ_C_* and *λ_R_* impacted the regrowth time. For example, for competitive interactions, combinations of values of *λ_C_* and *λ_R_* between 0.5-0.75 day^-1^ mm^-1^ generate regrowth times ranging 100-200 days (Fig 4b, sharp peaks), while those ranging between 0-0.5 day^-1^ mm^-1^ generate regrowth in about 50 days. Furthermore, for antagonistic interactions, when *λ_C_* < 0 < *λ_R_* regrowth is slower than when *λ_C_* > 0 > *λ_R_* by a factor of almost 2 (Fig 4a, 4d). These results suggest that the type of interaction changes the time of regrowth after radiotherapy and could contribute to variability in tumour responses.

**Figure 4.**
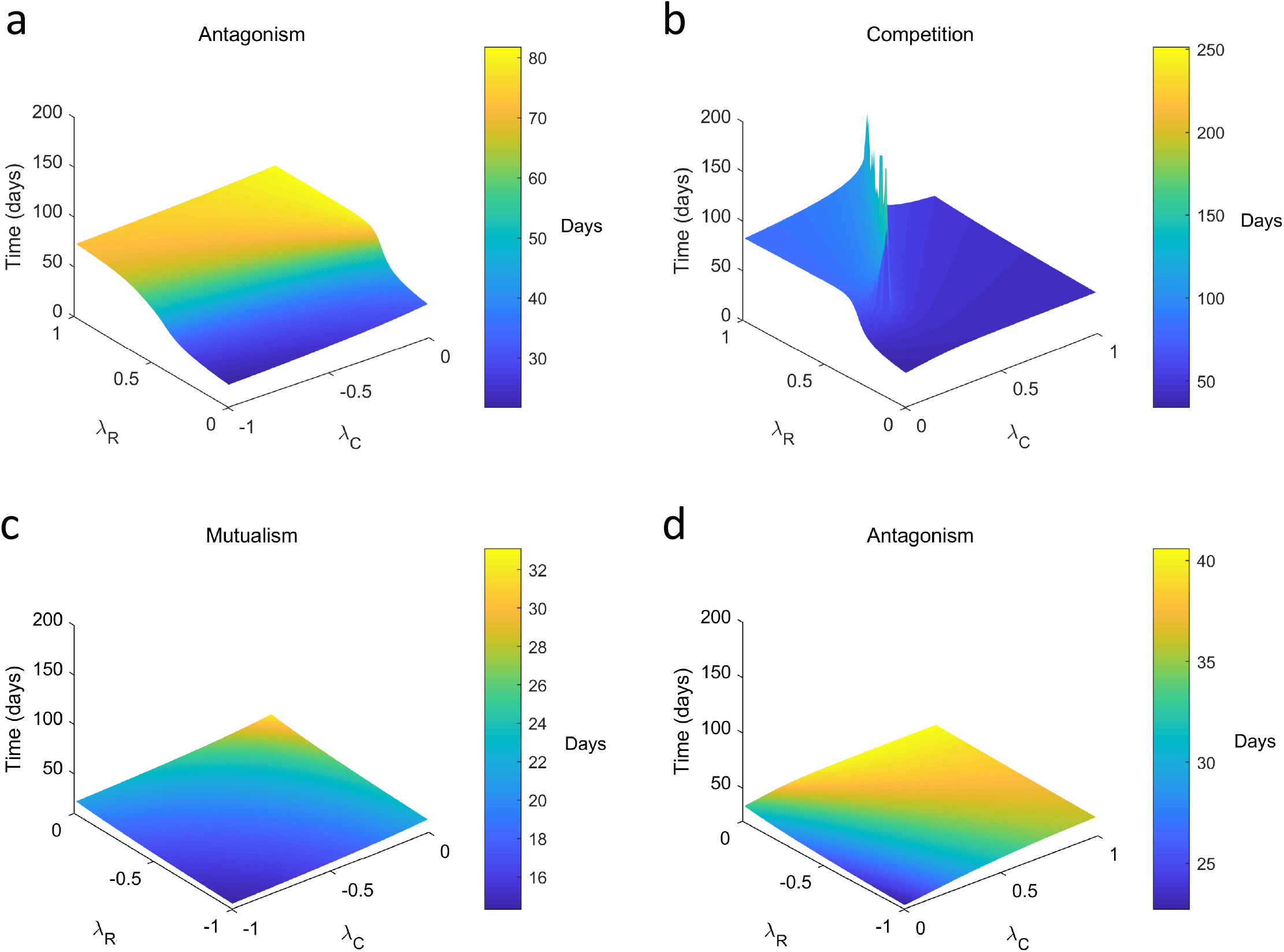
The nature of cellular interactions alters regrowth time after radiation, as determined by mathematical modelling. Mixed spheroids were grown in silico using a range of interaction parameters *λ_C_* and *λ_R_* until the volume reached 0.9 mm^3^. At this volume, spheroids were irradiated in silico and monitored for regrowth until they reached 0.9 mm^3^. Radiation damage was simulated using the linear quadratic model, for which input values (α and β) were determined from experimental PC3 clonogenic assays: ctrl cells, α = 0.44 ± 0.07 and β = 0.04 ± 0.01; res cells, α = 0.35 ± 0.06 and β = 0.03 ± 0.01. The numbers of control and resistant cells at the time of radiation depend on the values of *λ_C_* and *λ_R_,* whereas the numbers post radiation depend on the effect of radiation on each cell population, and on the values of *λ_C_* and *λ_R_*. See Supplementary Methods for more details on the mathematical model.

### Competition for oxygen between mixed populations determines cellular distribution

Indirect interactions between cells through competition for space and resources can influence the growth and response of spheroids to chemotherapy^33,34^. In our study, enhanced growth was observed in 3D but not in 2D. To identify potential environmental (i.e., indirect) mechanisms that contribute to the observed phenotype, we generated untreated homogeneous and mixed spheroids using fluorescent cells and evaluated the spatial distribution of each population using microscopy. Sections of mixed PC3 spheroids isolated on day 11 showed control cells located predominantly in the centre, a region that overlapped with staining for hypoxia, but not for proliferation (Fig. 5a). This localisation was detected in ~70 % of spheroids (SI Fig 8). The mixed and resistant spheroids also had reduced stromal staining and were visually less compact (Fig. 5a). No visual differences were observed in necrotic regions, or in spheroids isolated on day 5 (SI Fig 9), corroborating the measurements of proliferation and cell death ascertained by flow cytometry. Mixed DU145 spheroids also had a specific spatial structure, although inversely, with resistant cells in the centre and control cells in the periphery (SI Fig 10).

**Figure 5.**
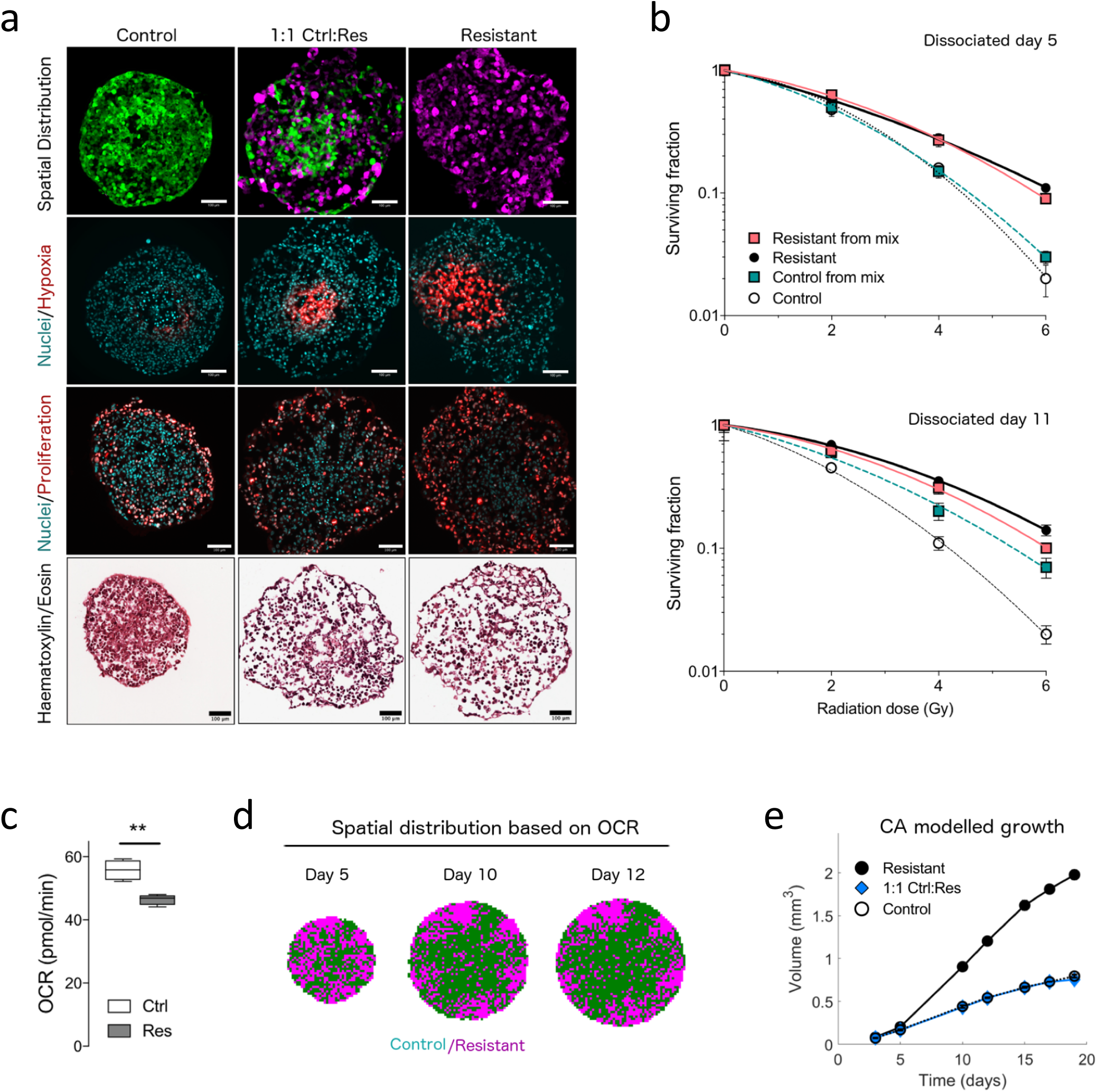
Indirect interactions through oxygen constraints influence cellular localisation of subpopulations in mixed spheroids. **(a)** Images showing spatial localisation of control (green) and resistant (magenta) cell populations, markers of hypoxia and proliferation, and H/E staining in homogeneous and mixed PC3 spheroids isolated on day 11. Scale bar, 100 μm. **(b)** Clonogenic survival of cell populations isolated from untreated spheroids dissociated at day 5 and day 11. **(c)** Oxygen consumption rate (OCR) of control and resistant PC3 cells. Values were normalized to cell density. **(d)** The spatial distribution of cell populations (control = green; resistant = magenta) based on OCR in mixed PC3 spheroids, as predicted by cellular automaton (CA) model. **(e)** Growth curves predicted solely by indirect interactions (i.e., competition for space and oxygen) in homogeneous spheroids (white and black circles) and mixed spheroids (blue circles). The CA model described in d was used to generate growth curves. See Supplementary Methods for more details. Data represent mean ± SD from 2 independent experiments for b and 4 independent experiments for c. * *P* < 0.05, ** *P* < 0.01, and *** *P* < 0.001, as determined by 1-way ANOVA (paired, one-tailed, alpha = 0.05).

We subsequently focused on interactions in mixed PC3 spheroids, which showed enhanced survival and resistance despite having competitive interactions. To determine whether localization of control PC3 cells in hypoxic regions resulted in enhanced radiation resistance, we dissociated spheroids at two time points, seeded them as single colonies and measured the clonogenic survival of each cell population. The radiation resistance of the control cells isolated from mixed PC3 spheroids increased 25% compared to controls isolated from homogeneous spheroids (radiation protection factor = 1.24, *P* = 0.04, Fig 5b). We verified that differences in intrinsic radiation sensitivity between control and resistant populations measured in normoxia were maintained under hypoxia, which could mask these differences (SI Fig 11).

Based on these results, we hypothesized that the spatial structure may result from competition for oxygen or differences in oxygen consumption rate (OCR) between the two cell populations. To test this hypothesis, we developed a cellular automaton (CA) model describing the changes in the size and structure of a 2D cross-section through a 3D tumour spheroid in which cells from control and resistant populations divide and die in response to the local oxygen gradient. In the model, cells consume oxygen supplied via diffusion, and mitosis occurs only if there is sufficient space around the dividing cell. Depending on the local oxygen concentration, cells can become hypoxic and enter cell cycle arrest or die via necrosis; if oxygen conditions improve, hypoxic cells can re-enter cell cycle. Parameter values for proliferation were derived from our flow cytometry data, while OCR values were measured using the Seahorse XF analyser and found to be 17% lower for resistant than for control cells (Fig 5c). The model was initialized by seeding equal numbers of control and resistant cells at day 0, based on the estimated values of *V_C_*(0) and *V_R_*(0) from the logistic model. On day 10, the resistant cells were located close to the spheroid periphery, while control cells were located close to the centre (Fig 5d). We then used the CA model to determine whether the growth curves of mixed spheroids could be explained by competition for oxygen. Although the CA model predicted the growth curves of the homogeneous cell lines (Fig. 5e), the model did not accurately capture the growth curves of the mixed spheroids under the assumption that the two cell populations compete for oxygen. Thus, although competition for oxygen may explain the spatial structure in mixed spheroids, the differences in OCR are not sufficient to explain the growth phenotype for PC3 spheroids.

We then investigated three known mechanisms by which hypoxic adaptation could enhance the survival and radiation resistance in control PC3 cells. Stabilization of the transcription factor HIF-1α under hypoxia can promote proliferation and survival^35^. However, basal levels and hypoxic induction of HIF-1α, as measured by Western blotting of PC3 cells incubated in normoxia/hypoxia for 24 hours, were not significantly different between control and resistant cells (SI Fig 12a). Another protein that can enhance cell survival in hypoxia is carbonic anhydrase 9 (CA9), which regulates intracellular pH^36^. CA9 expression was induced only in controls cells grown as monolayers under hypoxic conditions. However, CA9 expression, as assessed by fluorescence intensity in spheroid sections, was not significantly different *(P_adj_* = 0.07) among control, mixed, or resistant spheroids (SI Fig 12b). Furthermore, staining did not overlap exclusively with central regions of hypoxia in mixed spheroids, where control cells were found (SI Fig 12b). Instead, staining was observed both in peripheral and central regions, suggesting hypoxic-independent regulation of CA9 in 3D. We also investigated whether lipid droplet formation could enable survival in hypoxia^37^. However, no significant differences were found between the formation of lipid droplets in control spheroids and control cells isolated from mixed spheroids, as assessed by microscopy (SI Fig 12c) and flow cytometry (SI Fig 12d).

### Transferred factors from resistant cells enhance survival of controls in hypoxia and target the interferon pathway

We subsequently looked for evidence of direct interactions (i.e., cell-cell communication) that could explain the enhanced growth phenotype that we observed in untreated spheroids. Given that hypoxia was an important factor, we measured the growth of monolayer cells cultured in hypoxia (0.1% O2). Resistant and mixed populations grew significantly better in hypoxia than control cells (Fig 6a), in contrast to the phenotype measured in normoxia (Fig 1a). To evaluate whether this enhancement occurred through a transferred factor, we cultured cells in hypoxia for 120 h using Transwell inserts; in this system, cells can share factors (e.g., exosomes, microRNA, proteins) but are not in direct contact with each other. The number of control cells co-cultured with resistant cells was 48% higher than the number measured from homogeneous cultures. This effect was not observed in normoxia (Fig 6b). Altogether, our results indicate that both indirect interactions (through oxygen constraints), and direct interactions (through transfer of factor(s)) between mixed cell populations) enhance survival and radiation resistance in PC3 spheroids.

**Figure 6.**
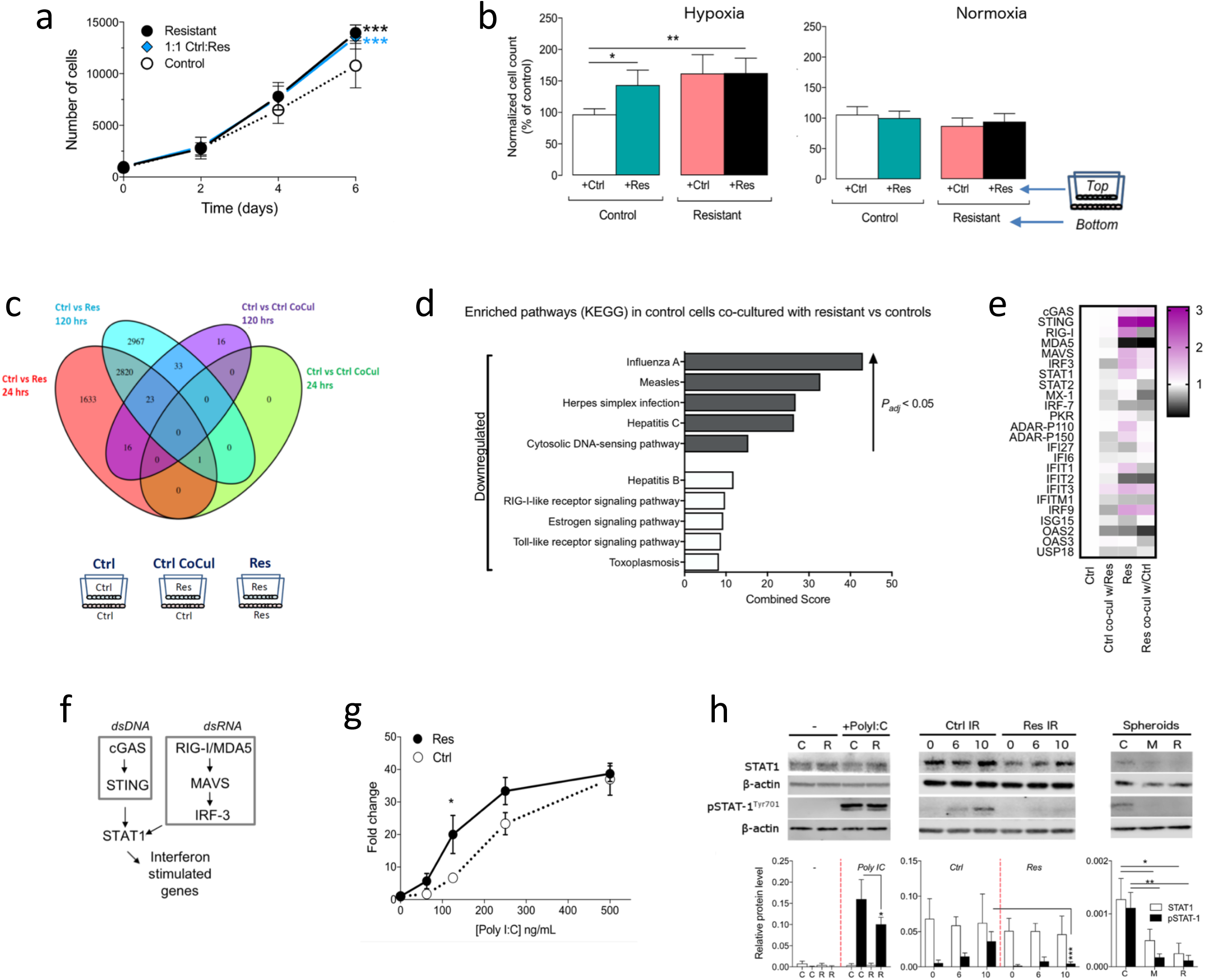
Altered interferon signalling correlates with hypoxic survival and radiation resistance. **(a)** Growth curves of PC3 cells grown as monolayers in hypoxia. Data represent mean ± SD from 4 independent experiments. **(b)** Cell count of control and resistant cells incubated separately or together in hypoxia (0.1% O2) and normoxia. The cells were cultured in Transwell inserts, in which cells share medium but are separated by a membrane. Data represent mean ± SD from 4 independent experiments. **(c)** Schematic illustrating set up of Transwell culture used for RNA sequencing (left side). Venn diagram of differentially expressed genes between control and resistant cells, and control and co-cultured controls, at 24 and 120 hours of hypoxia. Sequencing was performed on samples from 2 biological replicates. **(d)** Top 10 enriched KEGG pathways in control cells co-cultured with resistant cells in hypoxia. Gene enrichment analysis was performed using Enrichr^61^. Shaded bars indicate significant pathways (*P_adj_* < 0.05). **(e)** Expression of genes (fold change normalized to controls) from interferon pathway as assessed by qPCR from cells cultured in hypoxia 120 h. Data represent mean values from 3 independent experiments. **(f)** Simplified diagram of interferon pathway illustrating the convergence of the dsDNA and dsRNA pathways to activation of STAT-1 and interferon stimulated genes. **(g)** Activation of interferon pathway by control and resistant PC3 cells after exposure to synthetic dsRNA (poly I:C) for 24 hours in normoxia or hypoxia. Activation is measured by luminescence generated in HEK cells expressing a luciferase promoter coupled to an interferon response element promoter. Data represent mean ± SD from 3 independent experiments. **(h)** Western blots of STAT-1 and tyrosine701-phosphorylated-STAT1 expression in PC3 control and resistant cells after exposure to poly I:C (20 μg/mL) treatment for 6 h, to three different radiation doses (0, 6, and 10 Gy) after 48 h, or in untreated day 11 spheroids. Data represent mean ± SD from 4 independent experiments for poly I:C and radiation blots, and from 3 independent experiments for spheroid blots.* *P* < 0.05, ** *P* < 0.01, and *** *P* < 0.001, as determined by 2-way ANOVA followed by post-hoc t-tests with Holm-Sidak’s or Sidak’s correction for multiple testing.

We performed RNA sequencing to identify changes in the transcriptome of control cells co-cultured with resistant cells in hypoxia. We identified 88 genes that were differentially expressed in control cells co-cultured with resistant cells compared to controls alone at 120 h; no genes were differentially expressed between these two groups at 24 h (Fig 6c). Gene ontology analysis identified an enrichment of downregulated genes in viral pathways (measles, influenza, and hepatitis C), which are commonly activated in the interferon response (Fig 6d). An analysis of key genes in the interferon pathway using qPCR showed a small, but significant decrease in expression across the assayed genes (Fig 6e) in control cells co-cultured with resistant cells for 120 h compared to control cells in hypoxia (ANOVA_F(69,192) of interaction_ = 3.617, *P* < 0.001). In contrast, resistant cells had elevated levels of *STING* (*P_adj_* < 0.001), *DDX58* (*P_adj_* < 0.001), *MAVS* (*P_adj_* = 0.04), *IFIT3* (*P_adj_* < 0.01), and *IRF9* (*P_adj_* = 0.01), but lower levels of *MDA5* (*P_adj_* < 0.001) at 120 h hypoxia compared to control cells. These results are consistent with transfer of a factor (or factors) from resistant cells to controls that results in a downregulation of interferon pathway genes in control cells in co-culture, despite elevated levels in resistant cells under chronic hypoxia.

### Radiation resistant and mixed populations have altered interferon signalling

To determine whether control and resistant cell lines had basal differences in the interferon pathway, we measured the expression of genes involved in different branches of the pathway in control and resistant cells cultured in normoxia for 24 h (Fig 6f). Gene expression from the dsRNA and dsDNA branches were not significantly different between the two cell lines (ANOVAf(1,96) = 0.66, *P* = 0.41), except for *MDA-5* (*P_adj_* < 0.001), which was lower in resistant cells than in controls (SI Fig 13a). The two branches activate the phosphorylation of the transcription factor STAT-1 and subsequent transcription of interferon stimulated genes^38^. Phosphorylation of STAT-1 (pSTAT-1) was not different between either cell line (SI Fig 13b), and we measured no differences in the expression of downstream genes, except for a decrease in *MX-1* (*P_adj_* < 0.001) expression in resistant cells (SI Fig 13a). The two cell lines therefore do not substantially differ in the interferon pathway under basal conditions (i.e., 24 h normoxia).

Next, we investigated whether the two cell lines differed in their response to stimulation of the dsRNA and dsDNA sensors. To assess interferon induction to dsRNA stimuli, we exposed cells to the synthetic dsRNA polyI:C for 24 h, and transferred conditioned medium onto reporter HEK293 cells that luminesce when the interferon stimulated response element is activated. After exposure to 125 ng/mL of the synthetic dsRNA polyI:C for 24 h, resistant cells stimulated a 3fold higher response than control cells (*P_adj_* = 0.01); higher concentrations resulted in a common maximum plateau (Fig 6g). Stimulation with 20 μg of poly I:C for 6 h activated pSTAT-1 in both cell lines, but levels were ~ 40% lower (*P_adj_* = 0.01) in resistant cells than in controls (Fig 6h). However, no significant changes were measured between cell lines in expression of downstream ISGs with poly I:C stimulation, excepting for *MX-1* (*P_adj_* = 0.004, SI Fig 13c). Although the resistant cells are more sensitive to dsRNA stimuli in secreted interferon induction than controls, this production did not result in enhanced expression of downstream ISGs.

To assess response to dsDNA stimuli, we measured pSTAT-1 levels and expression of ISGs 48 h after 0, 6, and 10 Gy radiation. Induction of pSTAT-1 was observed in irradiated control cells, but not in irradiated resistant cells (Fig 6h). In downstream ISGs, expression of *DDX58, IFI27,* and *IFIT2* were significantly elevated in both cell lines after radiation (*P_adj_* < 0. 001), but expression of *IFITM1* was reduced by ~45% (*P_adj_* = 0.04) in irradiated resistant cells compared to irradiated controls (SI Fig 13d). We measured pSTAT-1 in untreated PC3 spheroids and found that it was induced in controls but not in mixed and resistant spheroids (Fig 6h). Taken together, these findings indicate that resistant cells respond differently to dsRNA and dsDNA stimulation, whereby the latter does not result in phosphorylation of STAT-1 at the measured time points and in 3D spheroids.

### Interactions between resistant and control cells determine cellular distribution in vivo, and their effects on growth and response are further influenced by angiogenesis

Our results demonstrate the importance of resistant and control cell interactions in an avascular 3D system. To determine whether the interactions could affect growth and response in vivo, where growth depends on nutrient and oxygen delivery through angiogenesis, we inoculated nude male mice with control, 1:1 mix, or resistant PC3 cells injected subcutaneously. Contrary to in vitro results, untreated control and mixed tumours grew faster than resistant tumours, reaching 200 mm^3^ an average of 10 days earlier than resistant tumours (Fig 7a).

**Figure 7.**
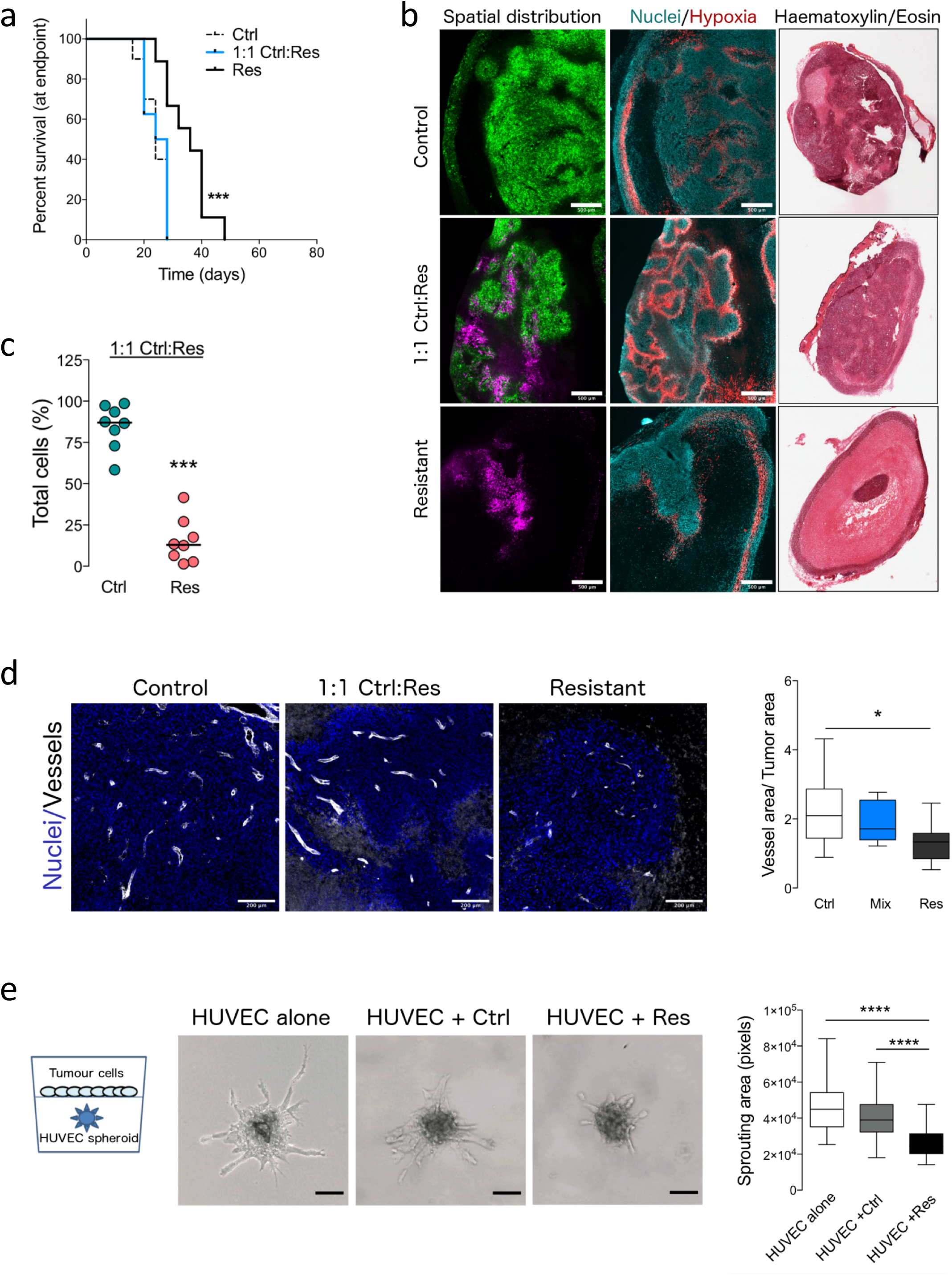
Cellular interactions alter spatial distribution in vivo and may be affected by angiogenesis. **(a)** Survival curves of untreated PC3 xenografts comprising control (n = 11 mice), 1:1 ctrl:res (n = 10 mice), or resistant cells (n = 11 mice). *** *P* < 0.001, as determined by Mantel-Cox test, followed by Bonferroni correction for multiple comparisons. **(b)** Representative images of whole tumour sections illustrating spatial distribution of control, resistant, and hypoxic cells in homogeneous and mixed xenografts. Scale bar, 500 μm. **(c)** Proportions of control and resistant cell populations isolated from mixed xenografts at end point (200 mm^3^), as measured by flow cytometry. (d) Representative images and quantification of vessel density in PC3 xenografts comprising control (n = 9 mice), mixed (n = 9 mice), or resistant (n = 10 mice) tumours. * *P_adj_* = 0.042, as determined by Kruskal-Wallis non-parametric test, followed by Dunn’s test for multiple comparisons. **(e)** Representative images and quantification of sprouting of endothelial spheroids cocultured with control or resistant cells in vitro, pooled from three independent batches of HUVEC endothelial cells (n = 20 spheroids/group/batch). Scale bar, 200 μm. **** *P* < 0.0001, as determined by Kruskal-Wallis non-parametric test, followed by Dunn’s test for multiple comparisons.

Survival outcome after 5 Gy radiotherapy was also not significantly different either among the three groups of PC3 tumours (control, 1:1 mix, resistant) (SI Fig 14a). However, the initial regrowth of mixed and resistant tumours after radiation was significantly higher *(P_adj_* = 0.04) than that of controls from days 28-40 (SI Fig 14b).

We then investigated whether interactions could affect the spatial distribution and survival of each population. In mixed tumours, control cells were evenly distributed, while resistant cells were localised in peripheral regions, where access to nutrients and oxygen would enable them to proliferate more rapidly (Fig 7b). Given that untreated mixed tumours comprised primarily of control cells (~80%) at the end point (Fig 7c), we reasoned that resistant cells could either die in vivo, rapidly proliferate and die, or stimulate the growth of control cells. Analysis using cell cycle distribution of single cells dissociated from 100% resistant tumours indicated a reduction in G0G1, and an increase in S compared to control tumours (SI Fig 14c), suggesting rapid proliferation and increased death. This was also confirmed by the large regions of necrosis in resistant tumours (Fig 7b).

Given the rate of cell turnover and the initial increase in regrowth after radiotherapy, we hypothesized that resistant cells had impaired/suppressed angiogenesis (and hence increased necrosis in the face of metabolic demands of proliferation). Staining for vessels using CD31 antibody showed that vessel density was < 31% (*P_adj_* = 0.042) in resistant tumours than in control tumours (Fig 7d). In vitro, HUVEC endothelial cells co-cultured in 3D with resistant cells sprouted 34% less than those co-cultured with control cells *(P_adj_* < 0.001, Fig 7e), corroborating the in vivo results. Together, these data suggest that while the cellular interactions between the populations do not measurably alter the endpoint measure (i.e., tumour volume) of growth or radiotherapy response, they affect the cellular localisation and survival of the populations, and substantially modify the microenvironment.

## DISCUSSION

### Role of cellular interactions on growth and radiation response

Our biological and mathematical results demonstrate that interactions between prostate tumour cell populations with different radiation sensitivities alter bulk tumour growth kinetics and radiation response of mixed populations. We mathematically predicted that both competitive and antagonistic interactions led to increased growth and resistance to radiation, and experimentally validated these predictions by showing that proliferation and survival rates of the control/resistant subpopulations were altered in mixed spheroids. Cellular interactions were mediated indirectly through oxygen constraints and directly through transferred factors. In vivo, these interactions did not measurably alter tumour growth kinetics or response to a single dose of radiation, but angiogenesis was a confounding factor. Nevertheless, cellular interactions affected the spatial structure and the immediate survival of cell populations after radiotherapy both in vitro and in vivo. Our study highlights that indirect and direct interactions between tumour cell populations with different radiation sensitivities could impact radiotherapy response in prostate cancer and thereby contribute to the observed heterogeneity in treatment response.

While antagonistic and mutualistic interactions have been shown to increase bulk tumour growth in human and mouse cancers^21–25^, we find here that competitive interactions can also have the same effect in prostate cancer. Furthermore, this competition did not result in the elimination of either clone, recapitulating the multiclonality observed in human prostate cancer ^4,5,9,39,40^. These findings are consistent with previous observations of competition between cell populations in fruit fly models ^41^, mammalian embryonic development^42,43^, and in breast cancer^24^. Collectively, these observations highlight that interactions between cell populations may have evolutionary advantages for cancer growth.

We also found that cellular interactions affect radiotherapy response when angiogenesis is not a confounding factor. This result was evidenced experimentally by the faster regrowth of mixed PC3 and DU145 spheroids after radiation and by enhanced survival rates of PC3 and DU145 control cells isolated from mixed spheroids after radiation than those isolated from homogeneous spheroids. However, overall survival was not different in vivo among irradiated control, mixed, and resistant tumours. Since resistant cells had higher rates of proliferation and impaired/suppressed angiogenesis both in vitro and in vivo, it is likely that high metabolic demands impaired tumour growth post-radiation. This hypothesis is consistent with the faster regrowth of resistant and mixed tumours in the initial days after radiotherapy. Differences in overall survival in vivo may therefore become apparent with higher doses of radiotherapy, which is known to impair angiogenesis ^44^ It remains unclear what differences in overall survival would be measured with fractionated radiotherapy, which has been shown to normalise vessels, improve blood flow, and increase oxygenation^45^.

Our mathematical simulations further demonstrated that the nature of the interaction (e.g., competition vs antagonism) affects the time of regrowth after radiation. After radiation, synthetic tumours with competitive interactions grew the slowest (50-250 days) while those with mutualistic interactions grew the fastest (16-32 days). Different values of interaction parameters within one type of interaction (e.g., competition) also had marked effects on tumour regrowth. These results suggest that it is not merely the nature of the interaction that matters, but also the extent of the effect exerted by each cell population on the other. While previous experimental and mathematical studies have shown that such cellular interactions increase the resistance of sensitive clones in mixed tumours to chemotherapy^19,20^, our experimental and mathematical data indicate that cellular interactions increase bulk tumour resistance to radiotherapy and increase the radiation resistance of sensitive clones. Future preclinical and mathematical studies could test whether accounting for these factors in the radiotherapy planning process improves tumour response to radiation and prevents recurrence.

### Impact of spatial distribution of tumour clones on radiotherapy outcome

Mechanistically, we found that the cellular interactions in PC3 spheroids result from oxygen constraints, and that these effects in turn dictate the spatial distribution of prostate cell populations in vitro and in vivo. This finding was supported both by experimental and mathematical models, where localisation of mixed populations was altered only in models where oxygen was a limiting factor (i.e., 3D spheroids, xenografts). Our results are consistent with those reported by Ibrahim-Hashim and colleagues, who found that the metabolic demands of breast cancer cells affect spatial localisation^14^. In our study, we found control PC3 cells predominantly near hypoxic regions and resistant cells near the periphery, while the inverse occurred in DU145 spheroids. While the localisation of resistant/sensitive cells differs between the cell lines, the general pattern of distinct localisations suggests that spatial heterogeneity of clones may be an important factor in treatment outcome. Recent studies have shown that the efficacy of chemotherapy strongly depends on the spatial distribution of tumour clones^33,46^. In another study, the localisation of resistant cells in the periphery was correlated with an increase in metastatic and invasive behaviours following chemotherapy. We hypothesize that these distinct spatial localisations could also affect radiotherapy outcome: delivery of non-uniform doses of radiation based on the spatial expression of prostate cancer antigen improved the tumour control probability in prostate cancer^47^.

### Impact of cell-cell communication on growth and radiation response

Direct interactions mediated by transferred factors from PC3 resistant populations enhanced the survival of controls in hypoxia. Control cells co-cultured with resistant cells in chronic hypoxia had a downregulation of interferon gene expression compared to control cells alone, suggesting that the transferred factor(s) downregulate interferon gene expression. Since proteomics analysis of conditioned medium requires no serum conditions (which makes cell survival difficult in chronic hypoxia), we were not able to determine the transferred factor shared between the populations. However, one possible factor could be an interferon. When stimulated with radiation, resistant cells induced several interferon genes (e.g., *DDX58, IFI27),* but did not phosphorylate STAT-1. Previous work confirms that cells resistant to DNA damage induce ISGs (e.g., *DDX58, IFI27)* independently of pSTAT-701^48^ and that they secrete interferons^49^. Furthermore, interferons can suppress angiogenesis^50^, consistent with reduced angiogenesis in the presence of resistant cells. However, qPCR data in control cells co-cultured with resistant cells in chronic hypoxia show downregulation in ISGs, inconsistent with the above explanations. Nevertheless, reduced interferon induction is potentially an important mechanism of adaptation in that it would protect against immune response after radiation^51^. Although interferons may be transferred from resistant to control cells, hypoxia is a confounding factor that downregulates interferon gene expression in cancer cells^51^. Together, these results highlight the importance of the interferon pathway in radiation resistance and merit further investigation, given the increased use of radiotherapy with immunotherapy (to capitalize on the immune system’s response to enhanced interferon signalling induced by radiation)^52^.

### Characterization of cellular interactions using mathematical modelling and cell biology

Experimental approaches to studying cellular interactions can be complex and time-consuming. The iterative and multidisciplinary approach applied here uses mathematical modelling to guide experimental design (and vice versa). Unlike approaches that primarily use experimental data to validate the predictions of math models, this iterative approach has two advantages. First, models can be used to test biological hypotheses and, in doing so, may generate new and unexpected findings. For example, we developed mathematical models built on experimental data to determine whether indirect and/or direct interactions contribute to an enhanced growth phenotype in mixed spheroids. This led us to discover that resistant cells enhance survival of controls through transferred factor(s) in mixed tumour spheroids. Second, simulations can be used to design concrete experimental protocols that can be used to accurately infer parameters in the theoretical models. In our study, parameter estimates were difficult to obtain with a small confidence interval in biological systems with > 20% noise. To overcome this issue, we ran simulations to determine the time points of collection for flow cytometry, which consequently enabled us to estimate interaction parameters reliably with narrower confidence intervals.

Although our mathematical model is relatively simple, it has three advantages over previous approaches. First, unlike other models used to study cellular interactions, it can describe a range of interactions, including competition, mutualism, and antagonism. Second, although the model does not account for different cell states (e.g., hypoxia, proliferation, necrotic) or for changes in interactions with time, it accurately predicted the major interactions occurring within spheroids. Third, the small number of parameters in the mathematical model can also be easily validated with experimental data. One limitation is that our model did not account for possible changes in the cellular interactions over time. Since tumour clonality can change with time, future studies are needed to investigate whether changing interactions (especially after treatment) could affect regrowth. Another limitation of the current model is that it does not account for angiogenesis, which seems to be a major confounding factor in vivo. We are now extending our Lotka-Volterra and CA models to account for the role of the vasculature in response to radiotherapy, exploiting existing models of vascular tumour growth where possible ^53–55^.

## CONCLUSIONS

In summary, we integrated experimental and computational models to investigate how tumour cells communicate with each other and their microenvironment to regulate tumour growth and response to radiation in prostate cancer. Our results suggest that indirect cellular interactions, their nature, and the spatial localisation of clones affect growth and radiation outcome. We also found that direct communication between tumour cell populations enhanced survival of control cells. Finally, in vivo analysis showed modification of the tumour microenvironment involving angiogenesis. Future studies will be needed to determine how to identify the type of interaction between tumour clones, the role of these interactions with fractionated therapy, and how these cellular interactions may be targeted to improve radiotherapy outcome in prostate cancer. Targeting higher doses to hypoxic areas may not be the best strategy to reducing resistance. Instead, mechanistic approaches targeting interferon signalling and reducing proliferation through metabolic inhibitors to prevent necrosis and hypoxia should be considered.

## METHODS

### Biological experiments

#### Cell culture

Two pairs of prostate cell lines, each containing cell populations with different intrinsic radiation sensitivities, were obtained from the Liu lab (Stonybrook, Canada): PC3 control and resistant^30^, DU145 control and resistant^31^. Three different batches of HUVEC cells, each containing 3 donors, were obtained from John Radcliffe Hospital and frozen in aliquots for onetime use. PC3 cells were cultured for up to 10 passages in DMEM (low glucose, pyruvate, GlutaMAX, Gibco) supplemented with 25 mM HEPES (Gibco), 10% foetal bovine serum (Sigma or Pan-Biotech), while HUVEC cells were cultured for up to 6 passages in EGM-2 media (Bulletkit, Lonza) comprising all components. The HEK293 interferon stimulated response element luciferase cell line were obtained from the Rehwinkel lab (Oxford, UK)^56^. Prostate cell lines were authenticated using STR analysis (Eurofins). All cells were maintained in an incubator (37 C, 5% CO2) and checked routinely for mycoplasma using MycoAlert Mycoplasma Detection Kit (Lonza).

#### Spheroid culture

Prostate spheroids containing homogeneous and mixed populations were generated from different ratios of control and resistant cell lines. Single cells were seeded (2×10^3^ cells/well) on ice, with 6-12 replicates per condition, in 96-well ultra-low attachment plates with round bottoms (Corning). Matrigel (Corning) was added (5% v/v) to medium (100 μL) to promote spheroid formation. Plates were centrifuged at 300 x g for 10 min at 4 °C. Fresh medium was added 3 days later (200 μL/well total volume). Half the medium was replenished every alternate day. HUVEC spheroids were generated using the hanging drop method as previously described^57^.

#### Growth assays

Proliferation assays were performed to determine the growth of homogeneous and mixed populations grown as monolayers and spheroids. For monolayer assays, single cells (1×10^3^ cells/well, 200 μL medium) were seeded in flat-bottom 96-well plates (Greiner), with 6 replicates per condition per experiment, and allowed to attach overnight. Medium was changed every 2 days. On days 0, 2, 4, and 6, cells were fixed using 4% paraformaldehyde (PFA), stained with Hoechst 33442 (0.1 mg/mL) for 10 min, washed in PBS, and scanned using the Celigo Cytometer (Nexelcom) to determine the cell number per well. For spheroids, homogeneous and mixed spheroids were generated and imaged repeatedly over time (2-3 times per week) using brightfield microscopy (4x objective; 0.3 NA, Leica DM IRBE, Hamamatsu). Spheroid volumes were calculated using SpheroidSizer^58^.

#### Generation of fluorescent cell lines

Prostate cell lines were transduced using lentiviral particles to produce stable, fluorescent cell lines. HIV lentiviral particles were generated by transfection of 293T cells with the vectors, pCDH1-CMV-GFP-EF1-Hygro or pCHD1-CMV-DsRed-EF1-Hygro (Systems Biosciences), and added to prostate cells. Twenty-four hours after transduction, particles were removed and medium containing hygromycin (Invitrogen) was added for selection (200 μg/mL for PC3 and 250 μg/mL for DU145). GFP- and DsRed-positive cells (top 30%) were sorted by flow cytometry 7 days after transduction, expanded for 7 days under selection, and then re-sorted (top 30%) by flow cytometry to obtain the brightest cells for subsequent experiments.

#### Flow cytometry of cell proportions, survival, and cell cycle

To determine proportions of cells in mixed spheroids, fluorescent cells were used to generate spheroids, dissociated at various time points, and analysed using flow cytometry. For each experiment, technical replicates comprised 6-8 pooled spheroids each. Spheroids were dissociated using 100 μL Accumax (Millipore) for 20 minutes at 37 °C. Single cells were washed with PBS, centrifuged at 300xg for 5 min, incubated with efluor-780 (1 μL/mL PBS; ThermoFisher Scientific) for 30 min on ice in the dark to distinguish live/dead cells, washed in PBS, and fixed for 10 min in IC Fixation Buffer (ThermoFisher Scientific). Samples were run on the LSR-II Fortessa X-20 using the 488, 561, and 633 lasers.

For cell cycle analysis, spheroids were incubated with EdU (10 μM final concentration) 12 h prior to dissociation, stained for live/dead cells, and fixed as described above. Cells were permeabilized and stained with Click-iT EdU Alexa Fluor 647 according to manufacturer’s instructions (ThermoFisher Scientific). After washing in 1X saponin, cells were incubated 30 min with FxCycle Violet Stain (1:1000 stock diluted in 300 μL of 1X saponin; ThermoFisher Scientific) before being analysed on the Attune NxT Flow Cytometer (ThermoFisher Scientific) using the 405, 488, 561, and 633 lasers. Data were analysed using FlowJo (Treestar, Inc).

#### Radiation response of individual populations and bulk spheroids

Survival of cell lines after radiation was measured using a clonogenic assay^59^. Briefly, cells in the exponential growth phase were seeded in six-well plates and irradiated at a range of doses (0 to 6 Gy) using a Cs-137 irradiator (dose rate 0.89 Gy/min). For clonogenic assays performed under hypoxia, cells were seeded and allowed to attach for 3 hrs, placed in a hypoxia chamber (InVivo Chamber 300) for 6 hours at 5% CO2 and 0.1% O2, and then placed in sealed chambers for irradiation. After 10-14 days incubation, surviving colonies were stained with crystal violet and counted. The surviving fraction was calculated as: (number of counted colonies/number of seeded cells) x plating efficiency.

To determine whether bulk radiation response was altered in mixed populations, PC3 cells were seeded as spheroids (n = 15 per dose per group) with 4 groups: ctrl, 9:1 ctrl:res, 1:1 ctrl:res, and res. After formation, spheroids were irradiated on day 3 using a range of doses (0, 2.5, 5, 7.5, 10, 15, and 20 Gy) and imaged for up to 60 days to monitor regrowth. Spheroids that reached 3 times the initial starting volume (i.e., the volume measured on day 3) were considered to have regrown. The experiment was repeated on a smaller scale using single doses of radiation on ctrl, mixed, and resistant spheroids (n = 20/ group) from both PC3 and DU145 cell lines. To measure changes in the radiation response of PC3 cell populations within spheroids, untreated homogeneous and mixed spheroids were grown until day 5 or 11, dissociated using Accumax, seeded as single cells for clonogenic experiments, and allowed to attach for 6 hours prior to radiation. Fluorescent colonies were stained with Hoechst and counted using Celigo.

#### Staining and microscopy

To investigate the spatial distribution of cell populations, hypoxia, proteins (Ki67, Ca9), and necrosis, spheroids and xenografts were fixed as before^60^ and sectioned. For spatial distribution of fluorescent populations, sections were hydrated in PBS, stained for 10 min with Hoechst (1 μg/mL in PBS, Sigma) to visualize nuclei, and mounted using ProLong Diamond Antifade Mountant (ThermoFisher). For hypoxia, spheroids and xenografts pre-treated with the hypoxia drug EF5 (University of Pennsylvania) were stained using anti-EF5 antibody as previously described^60^. For Ki67 and Ca9, spheroid sections were permeabilized for 10 min using PBS containing 0.3% Tween-20. After permeabilization, sections were washed 3×5 min in PBS, blocked using 5% goat serum in PBS containing 0.1% Tween-20, and incubated overnight at 4 °C with primary antibody. Sections were then washed 3 x 5 min in PBS, incubated for 1 h with goat anti-rabbit Alexa Fluor 647 (4 μg/mL, ThermoFisher), washed, and stained with Hoechst 33342 (50 μg/mL, Sigma) for 10 min. After a final wash, slides were mounted and imaged using epifluorescence microscopy (20x objective, 0.30 NA, 0.64 μm resolution, Nikon Ti-E). Primary antibodies were: Ki67 (clone SP6, 1:100, Vector Laboratories), and Ca9 (clone M75, 2.5 μg/mL, Bioscience Slovakia). Sections were stained using H*E and imaged using a Bright Field Slide Scanner (Aperio) to visualize necrosis.

#### Oxygen consumption measurements

Cells (1.2×10^4^/well) were seeded using normal medium in Seahorse XF 96-well microplates (Agilent) and allowed to attach overnight. Prior to the assay, cells were washed with and incubated in assay medium (200 μL) for 2 hours at 37 °C without CO2 to degas the medium. Oxygen consumption rates (OCR) were measured from each population using Seahorse XF Analyzer (Agilent Biosciences). After OCR measurements were obtained, cells were fixed in 4% PFA, stained using Hoechst, and counted using Celigo.

#### Western blotting

Cells (2.0×10^5^/well) were seeded in 6-well plates (Corning) and allowed to attach overnight. Once the medium was changed, cells were treated with polyinosinic-polycytidylic acid (Poly I:C, 20 μg/mL, Sigma-Aldrich) for 6 h, incubated in hypoxia (0.1% O2) for 24 h, or incubated for 48 h after radiation (0, 6, 10 Gy). Cells were washed with cold-PBS and lysed on ice using RIPA buffer (100 μL/well, Sigma) containing freshly added phosphatase (phosSTOP, Roche) and protease inhibitors (cOmplete EDTA-free, Roche), as per manufacturer’s instructions to collect supernatants. For HIF1-α expression, cells were lysed in the hypoxia chamber using 9M urea containing β-mercaptoethanol, sonicated using a water bath, and centrifuged.

After electrophoresis of samples (4-12% NuPage Tris-Bis gels, ThermoFisher) and semidry transfer, membranes (PDVF, Immobilon-FL, Millipore) were blocked for 1 h using diluted Odyssey buffer (1:1 in TBS, LI-COR Biosciences), and probed for expression of proteins following manufacturer’s instructions. Images were acquired and quantified using Odyssey Fc (LI-COR Biosciences). Primary antibodies were: HIF1α (clone 54, 1:500, BD Biosciences); CA9 (clone M75, 1:1000, Bioscience Slovakia); p-STAT1-Tyr701 (9167, 1:1000, Cell Signalling); STAT-1 (9172, 1:1000, Cell Signalling); β-actin (4967, 1:5000, Cell Signalling). Secondary antibodies were: goat anti-rabbit IgG or goat anti-mouse IgG (IRDye 680RD or 800CW, 1:10000, Odyssey, LICOR).

#### co-culture in hypoxia

Co-culture experiments were performed to measure whether factors transferred from resistant cells enhanced survival. Cells were seeded (3.0×10^4^/bottom well and 1.0×10^4^/insert) in 12-well plates and in Transwell inserts, and allowed to attach overnight. Control and resistant cells were cultured in the 12-well plates, as well as in the Transwell inserts, as follows: ctrl/ctrl, ctrl/res, res/ctrl, res/res. Once medium was changed, the plates were placed into normoxia or hypoxia (0.1% O2) for 24 h and 120 h. Cells were fixed using 4% PFA, stained with Hoechst (50 μg/mL), and counted using Celigo.

#### Poly I:C transfection for Western blotting

To induce an interferon response through the dsRNA pathway, cells were seeded in 6-well plates (2×10^5^ cells/well), allowed to attach overnight, and transfected using poly I:C. Transfection was performed for 6 h using OPTI-MEM GlutaMax (Gibco) containing poly I:C (0 and 20 μg/mL) and lipofectamine 2000 (5 μL) following manufacturer’s instructions.

#### Luciferase reporter assay

Differences in the functional activation of the interferon pathway between cell lines was measured using a luciferase reporter assay. On day 1, PC3 control and resistant cells (1×10^4^/well; 100 μL) were seeded in 96-well plates. On day 2, PC3 cells were transfected with lipofectamine (1 μL/well) and poly I:C (0-500 ng/mL, 100 μL total volume/well) for 1 h, washed 3 times with PBS, and incubated in normal medium. Cells from the recombinant HEK293 cell line, stably expressing the firefly luciferase gene under the control of interferon-stimulated response element^56^, were seeded (2×10^4^/well; 100 μL) in 96-well plates and allowed to attach. On day 3, the medium of the HEK293 cells was replaced with conditioned medium (50 μL) from PC3 cells for 24 h. On day 4, One-Glo Luciferase reagent (50 μL/well) was added to HEK293 cells to lyse cells before measurement of luminescence.

#### RNA sequencing

Bulk RNA sequencing was performed to measure changes in gene expression of cells cocultured in hypoxia. Co-culture experiments were set up as described above. Total RNA was extracted at 24 h and 120 h using mirVana Isolation Kit according to manufacturer’s instructions and quantified using TapeStation (Agilent Biotechnologies). Bulk RNA sequencing was performed on biological duplicates using Illumina sequencing (75 million reads). Differences in gene expression were quantified using the DE-Seq2 package in R (R v 3.4) and analysed for statistical significance using adjusted P-values with a false-discovery rate of 0.1. Gene enrichment was analysed using the KEGG pathway in Enrichr^61^.

#### RT-PCR and qPCR

Differences in expression of several genes in the interferon pathway were measured using qPCR. Cells (2.0×10^5^/well) were seeded in 6-well plates (Corning), allowed to attach overnight, and treated with normoxia or hypoxia for 24 h. Total RNA was extracted using TRI-reagent (Sigma), isopropanol (Sigma), and 1-bromo-3-chloropropane (Sigma), and 1 μg was reverse transcribed into cDNA using High Capacity cDNA Reverse Transcription Kit (Thermo Fisher). Each sample was amplified in triplicate using custom-designed primers (Sigma, Supplementary Table 2) in a HT7900 Real Time PCR System using SensiMix SYBR-Green Mix (Bioline) with standard cycling conditions: 10 min at 95°C, 15 sec at 95°C (40 cycles), and 45 sec at 60°C. Gene expression was analysed with the 2^-ΔΔCT^ method using *RPL11* as the reference gene.

#### Xenograft tumour growth and radiation response

To determine whether interactions affected tumour growth kinetics in vivo, we measured growth curves of xenografts from three groups: control, 1: 1 mix, or resistant. Fluorescent PC3 cells (5 x 10^6^ cells/100μL comprising PBS and Matrigel, 1:1) were inoculated subcutaneously in the right flank of 36 male nude mice (nu/nu, 5-6 weeks, 27.1 ± 2.73 g, Envigo) with 12 mice/group. Sample sizes were estimated from a pilot study (n = 10 mice) and from previous work^30^. Once tumours reached 70 mm^3^ (calculated L×W×H×π/6), they were measured with callipers every 3 days until they reached 200 mm^3^. To establish whether interactions affected radiation response in vivo, 36 male mice were inoculated on the right flank with control, 1: 1 mix, or resistant PC3 cells. When tumours reached 70 mm^3^, they were irradiated with a single dose of 5 Gy using a Gulmay 320 kV X-irradiator (2.0 Gy/min) and measured every 3 days for 72 days. Animals that did not develop tumours within 4 weeks of inoculation were excluded from experiments. However, animals with regressing tumours post-radiation were included in survival analysis and marked as “censored” events. Randomization was not used post-inoculation, but a partial-blinding system (cage and ear tag labels) was used to track tumour growth and label ex vivo samples. All animal procedures were conducted in accordance with the UK Animal Scientific Procedures Act of 1986 (Project License Number PCDCAFDE0) and were approved by the local ethics committee at the University of Oxford.

For spatial distribution and flow cytometry analysis in untreated xenografts, mice were injected with EF5 (10 mM, saline) and with EdU (10 mM, PBS) 2 h prior to being culled. Parts of the tumour were immediately dissected for analysis using flow cytometry, while other parts were fixed in 4% PFA for immunofluorescence staining. For flow cytometry, tumours were minced and digested in Hank’s Buffered Saline Solution (ThermoFisher) containing collagenase II (2 mg/mL, Worthington) and DNAse I (2 U/mL, ThermoFisher) for 30 min at 37 °C with shaking. Samples were put through a cell strainer (50 μm, Sysmex) and washed with ice-cold PBS. After centrifugation (300xg, 5 min, 4 °C), single cells were processed for flow cytometry of survival and cell cycle as described above. Immunofluorescence staining was performed as describe above for spheroid sections.

#### Endothelial sprouting assay

To investigate whether control and resistant tumour cells affected angiogenesis, we measured sprouting of HUVEC spheroids in the presence of tumour cells. HUVEC spheroids were embedded in collagen gel in 24-well plates and incubated 1 h to allow the gel to solidify.^57^ Tumour cells (2×10^4^) cells were placed on the top of the gel in EGM-2 medium and incubated for 24 h. For each spheroid, a single image was acquired at the z-plane with the largest diameter in focus (Evos). The area of sprouting from this single image was measured using ImageJ.

### Mathematical modelling

#### Non-spatial mathematical models

We used the logistic population growth model to describe the growth of homogeneous tumour spheroids^62^. According to this model the rate of change of spheroid volume *V*at time *t* is given by

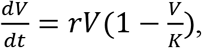

where *r*>0 represents the growth rate, *K*>0 is the carrying capacity (the limiting volume of the spheroid) and *V*(*t*=0) = *V*_0_ where *V*_0_ is the spheroid volume at *t*=0. The analytical solution to the logistic model is given by

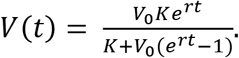

The growth of and interactions between the control and resistant populations were described by a Lotka-Volterra-type model given by

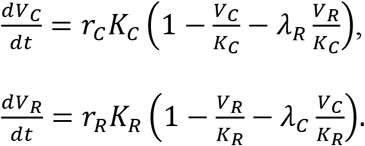

In the model above, *V_C_* and *V_R_* represent respectively the volumes of control and resistant populations, *r_C_* and *r_R_* their initial growth rates, *K_C_* and *K_R_* carrying capacities, and *V_C_*(0) and *V_R_*(0) their initial volumes. The parameters *λ_C_* and *λ_R_* describe the magnitude of the effect that control cells have on resistant cells, and resistant cells have on control cells, respectively. Typically, the signs of the interaction parameters are constrained so that the model can only describe either competition (*λ_C_* >0 and *λ_R_* >0) or mutualism (*λ_C_* <0 and *λ_R_* < 0)^63^. We do not put such constraints on *λ_C_* and *λ_R_* which means that our model can describe most types of biological interactions found in ecology. Further details describing how the values of the model parameters were estimated are included in Supplementary Methods.

#### Spatial computational model

To gain further insight into the growth of heterogeneous spheroids we developed a spatially-resolved cellular automaton (CA) model. Our CA model replicated the changes in the size and structure of a 2D cross-section through a 3D tumour spheroid suspended in culture medium. The model couples a set of automaton elements arranged on a regular 2D grid to a partial differential equation (PDE) describing the distribution of a growth-rate-limiting nutrient (here, oxygen) which is supplied from the culture medium surrounding the spheroid. Our model simulations are initialised by placing a circular cluster of cells in the centre of the grid: this imitates seeding a spheroid in a Petri dish. The control and resistant cells differ in cell cycle times, oxygen consumption rates, sensitivity to changing oxygen concentration and rates of lysis. Both cell populations consume oxygen as it diffuses from the medium and divide. When local oxygen concentration drops below threshold values cells become hypoxic and eventually die via necrosis. Furthermore, cell division depends on local cell density. Further details pertaining to the model implementation are described in Supplementary Methods.

### Statistical Analysis

Data were evaluated for homogeneity of variance and for normality. For all biological assays unless indicated below, statistical significance was evaluated using 2-factor ANOVA followed by Bonferroni, Sidak, or Dunnett multiple-testing correction (α= 0.05). Effect sizes were approximated from pilot studies or from literature to ensure power (β = 0.8) for biological experiments. For clonogenic assays, the radiation protection factor (RPF) was calculated as the area under the dose-response curve (AUC) for the resistant cell lines divided by that of the control lines; AUC values were analysed for significance using a Student’s t-test (paired, one-tailed, α= 0.05). For spheroid control probability experiments, dose-regrowth curves were fitted and analysed by nonlinear regression using a log-sigmoidal model with variable slope (Prism 5.0, GraphPad). Values of spheroid control probability (SCP_50_) are reported as the dose at which 50% of spheroids did not regrow, along with the 95% CI of the SCP_50_. For additional regrowth experiments, survival curves were analysed using the Mantel-Cox (log-rank) test. Data represent mean ± SD from independent experiments, as indicated.

### Code and data availability

Code and data are available upon request. RNA sequencing data is available at the NCBI GEO database with accession number XXX.

## AUTHOR CONTRIBUTIONS

PK, ALH, M Partridge, and HB conceived the project. PK, M Paczkowski, LKS, ALH, M Partridge, and HB designed experiments. PK, M Paczkowski, and AM performed cell biology experiments. PK, JO, SL, JK, JC, and BM performed experiments relating to xenografts. M Paczkowski performed mathematical modelling. WWK performed bioinformatic analysis. PK, M Paczkowski, and WWK analysed data. PK, M Partridge, ALH, and HB supervised project. PK, M Paczkowski, ALH, and HB wrote the manuscript. All authors edited/reviewed the manuscript.

## ACKNOWLEDGMENTS

We thank Yunhong Cao and Karla Watson for technical assistance. This research was supported by the Medical Research Council, Cancer Research UK, and the Engineering Physical Sciences Research Council (grant numbers: C5255/A12678, C2522/A10339, and C56606/A21440). PK, JK, BM, and M Partridge were funded by the CRUK/EPSRC Oxford Cancer Imaging Centre. AM and ALH were funded by Breast Cancer Now (grant number: 2015MayPR479). WWK was funded by KTH Royal Institute of Technology. M Paczkowski was funded by EPSRC LSI grant.

