## Supplementary Information for "Radiation resistant cancer cells enhance the survival and resistance of sensitive cells in prostate spheroids"

**SUPPLEMENTARY TABLES**

**Supplementary Table 1.** Initial volumes of control and resistant cells in untreated homogeneous spheroids from PC3 and DU145 cell lines.

| Cell line | $V_C(0)$ | $V_R(0)$ |
|---|---|---|
| PC3 | 0.040 *(0.033, 0.047)* | 0.036 *(0.032, 0.042)* |
| DU145 | 0.030 *(0.025, 0.036)* | 0.046 *(0.041, 0.052)* |

Values indicate mean value and 95% confidence intervals. $V_C(0)$ = initial volume in control spheroids; $V_R(0)$ = initial volume in resistant spheroids;

**Supplementary Table 2.** Primer sequences for real-time PCR.

| Gene | Forward primer | Reverse primer |
|---|---|---|
| *ADAR-1 p150* | CTTCCAGTGCGGAGTAGCG | GTGACGGTGTCTGCTTTCCA |
| *ADAR-1 p110* | AAGGTCAGGAAGATTGGCGA | CTTGGTAAACGAACTTGGGCT |
| *cGAS* | TCCAAGAAGAAACATGGCGG | CACGCAGTTATCAAAGCAGAG |
| *DDX58* | CAAGCCTTCCAGGATTATATCCG | AGTCCAGAATAACCTGCATGGT |
| *IFIT1* | TACCTGGACAAGGTGGAGAA | GTGAGGACATGTTGGCTAGA |
| *IFIT27* | CTCCTTCTTTGGGTCTGGCT | GCCACAACTCCTCCAATCAC |
| *IFIT2* | TGTGCAACCTACTGGCCTAT | TTGCCAGTCCAGAGGTGAAT |
| *IFIT3* | CTTCAGTATTTACTTGAGGCAGAC | CTTGGTGACCTCACTCATGAC |
| *IFITM1* | AACCACACTTCTCAAACCTTCAC | CCTGTCCCTAGACTTCACGG |
| *IFI6* | TCTCCTCCAAGGTCTAGTGAC | TCTCCTCCAAGGTCTAGTGAC |
| *ISG15* | GCGAACTCATCTTTGCCAGTA | CCAGCATCTTCACCGTCAG |
| *IRF3* | CTCGTGATGGTCAAGGTTGTG | AGTTTATTGGTTGAGGTGGTGG |
| *IRF7* | AAGGGCTTCCCCTGACTG | TCTACTGCCCACCCGTACA |
| *IRF9* | AGAAAGTACCATCAAAGCGAC | TTGTGTCTGTAACTTCCTGTG |
| *ISG15* | CGAACTCATCTTTGCCAGTA | CCAGCATCTTCACCGTCAG |
| *MAVS* | GTACCCGAGTCTCGTTTCCT | ATGAAGTACTCCACCCAGCC |
| *MDA5* | TTAACAGGCTCTGATTGCTC | TCTCTTCATCTGAATCACTTCCC |
| *MX-1* | GTTACCAGGACTACGAGATTGAG | GATGAGTGTCTTGATCTTATACCC |
| *OAS2* | ATCTGTATAAATCCTCGGACCT | CCTTTCACACTCTTTGTACCA |
| *OAS3* | TACCAGCAGTGTACCAAGATCTC | TTTCAGGAAGTCTCCAACAGTC |
| *PKR* | ATCTGACTACCTGTCCTCTG | GAGACCATTCATAAGCAACGA |
| *RPL11* | GCAAACTCTGTTCAACATCTG | CATACTCCCGCACTTTAGAC |
| *STAT1* | TACACCTACGAACATGACCCT | TCACCAACAGTCTCAACTTCAC |
| *STAT2* | CCCGCTGACTGAAATCATCC | AGTTCATCCACCTGTCTATTAGAG |
| *STING* | GTTTGCCATGTCACAATACAGTC | TGCCCACAGTAACCTCTTCC |
| *USP18* | CAGGATATTGAAGAGGATCACGG | CTTAATCAGGTTCCAGAGTTTGAG |

#### SUPPLEMENTARY FIGURE LEGENDS

**Supplementary Figure 1.** Intrinsic radiation sensitivity between control and resistant populations from PC3 and DU145 cell lines in normoxia. Clonogenic survival of cells was measured 10-14 days after irradiation with 0, 2, 4, and 6 Gy. Data represent mean ± SD from 3 independent experiments. Statistical significance was evaluated using Student's t-test on area-under-the-curve values (paired, one-tailed, $\alpha$=0.05). RPF = radiation protection factor.

**Supplementary Figure 2.** Gating strategy for flow cytometry experiments measuring proportions of cell populations in spheroids. **(a)** After spheroids (seeded 1:1 ctrl:res) were dissociated, single cells were isolated from debris (SSC-A vs FSC-A) and doublets (FSC-H vs FSC-A). After gating of live and dead cells (efluor 780; Apc-Cy7-A vs FSC-A), control (GFP+) and resistant (DsRed+) cells were identified (Alexa Fluor 488-A vs PE-Texas Red-A). Percentages shown are derived from the proportion of cells of the previous gate. A total of 20,000 events were collected. SSC-A, side-scatter area; FSC-A, forward-scatter area; FSC-H, forward-scatter height. **(b)** Sample plots of dissociated spheroids from days 5, 10, and 15.

**Supplementary Figure 3.** Interaction parameters can be estimated mathematically using spheroid growth curves and proportions of cell populations. **(a)** Values of interaction parameters estimated using simulated growth curves alone. Values of interaction parameters, $\lambda_C$ and $\lambda_R$ ($day^{-1}$ $mm^{-1}$), estimated from 100 simulations of in silico growth curves of heterogeneous populations containing 1:1 mix of control: resistant cells. Growth curves were generated with different levels of noise to simulate biological variance. **(b)** Values of interaction parameters estimated using simulated growth curves and the proportion of each population. Data represent mean ± 95% confidence interval.

**Supplementary Figure 4.** Gating strategy for flow cytometry experiments measuring cell cycle for each cell population in homogeneous and mixed spheroids. After excluding debris (top left), doublets (top middle), and dead cells (top right), control (GFP+) and resistant (DsRed+) cells were analysed for cell cycle proportions. The uptake of the nucleoside analogue 5-ethynyl-2 deoxyuridine (EdU) was detected both in cells proliferating in S phase and in cells that previously took up EdU in S phase and then cycled into $G_0G_1$ during the incubation time. Gating was thus performed to include all proliferation (EdU+ cells) to account for turnover. Percentages shown are derived from the proportion of cells of the previous gate, but were recalculated for cell cycle analysis to obtain proportions from only the selected gates.

**Supplementary Figure 5.** Interactions between control and resistant populations alter cell death in untreated mixed spheroids, as assessed by flow cytometry. Single cells were isolated from spheroids seeded with 100% control cells, 100% resistant cells, or 1:1 mixture of ctrl:res (top panel). Percent of dead cells in PC3 spheroids (middle panel) and DU145 spheroids (lower panel) measured over time. Data represent mean ± SD from 4 independent experiments. ** $P < 0.01$, *** $P < 0.001$, as determined by 2-way ANOVA followed by Bonferroni correction for multiple testing.

**Supplementary Figure 6.** Interactions between control and resistant populations alter cell death in irradiated mixed spheroids, as assessed by flow cytometry. Single cells were isolated from irradiated spheroids seeded with 100% control cells, 100% resistant cells, or 1:1 mixture of ctrl:res (top panel). Percent of dead cells in PC3 spheroids (middle panel) and DU145 spheroids (lower panel) measured over time. Data represent mean ± SD from 4 independent experiments. ** $P < 0.01$, *** $P < 0.001$, as determined by 2-way ANOVA followed by Bonferroni correction for multiple testing.

**Supplementary Figure 7.** Interactions between control and resistant populations alter cell cycle in irradiated mixed spheroids, as assessed by flow cytometry. Cell cycle of control and resistant cell populations isolated from mixed and homogeneous spheroids from both PC3 and DU145 cell lines. Data represent mean ± SD from 4 independent experiments. * $P < 0.05$, ** $P < 0.01$, *** $P < 0.001$, as determined by 2-way ANOVA followed by Bonferroni correction for multiple testing.

**Supplementary Figure 8.** Images showing spatial localisation of control (green) and resistant (magenta) cell populations in 16 different mixed PC3 spheroids isolated on day 11. Scale bar = 100 μm.

**Supplementary Figure 9.** Images showing spatial localisation of control (green) and resistant (magenta) cell populations, markers of hypoxia (EF5, red), proliferation (Ki67, red), nuclei (Hoechst 33342, cyan), and H/E staining in homogeneous and mixed PC3 spheroids isolated on day 5. Scale bar = 100 μm.

**Supplementary Figure 10.** Images showing spatial localisation of control (green) and resistant (magenta) cell populations, and H/E staining in homogeneous and mixed DU145 spheroids isolated on day 5. Scale bar = 100 μm.

**Supplementary Figure 11.** Intrinsic radiation sensitivity between control and resistant populations from PC3 and DU145 cell lines maintained in hypoxia. Clonogenic survival of cells was measured 10-14 days after irradiation in hypoxia (0.1% $O_2$). Data represent mean ± SD from 3 independent experiments. Statistical significance was evaluated using Student's t-test on area-under-the-curve values (paired, one-tailed, $\alpha$=0.05).

**Supplementary Figure 12.** Common mechanisms of hypoxic adaptation do not account for enhanced survival and radiation resistance in mixed PC3 spheroids. **(a)** Protein levels of hypoxia inducible factor 1$\alpha$ (HIF-1$\alpha$) and carbonic anhydrase 9 (CA9) measured from whole lysates of PC3 control (C) and resistant (R) cells treated with normoxia (20% $O_2$), 100 μM deferoxamine (DFO), or hypoxia (0.1% $O_2$) for 24 hours. Protein densities were normalised to β-actin values (mean ± SD, n = 3 independent experiments, $P_{adj} < 0.001$, 2-way ANOVA with Bonferroni correction). **(b)** Representative images of staining for Ca9 (green) and hypoxia (EF5, magenta) in control, mixed, and resistant spheroids. Scale bar, 100 μm. Quantification of mean fluorescence intensities of CA9 staining (n = 8 spheroids/group, $P$ = 0.07, Kruskal-Wallis test). **(c)** Representative images of staining for control cells (GFP+, green), resistant cells (DsRed+, magenta), and lipids (LipidTox 647, yellow) in mixed spheroids. Scale bar for left and middle panels, 200 μm. Scale bar for right panel, 100 μm. **(d)** Flow cytometry measurement of lipid droplet intensity in control, mixed, and resistant spheroids dissociated at day 11. Gating strategy for flow cytometry is shown on left-hand side: after exclusion of debris and doublets, live cells (RL3-A) were selected to separate control (BL1-A) and resistant (YL1-A) populations. Median fluorescent intensity of lipid staining (RL1-A) was quantified for each cell population, as shown on the right-hand side. Data represent mean ± SD (n = 3 independent experiments, * $P_{adj} < 0.05$, 1-way ANOVA with Tukey correction).

**Supplementary Figure 13.** Expression of interferon pathway genes in PC3 control and resistant cells under basal and stimulated conditions, as assessed by qPCR. **(a)** Expression of genes under baseline conditions (24 h normoxia). **(b)** Representative Western blot of STAT-1 and phosphorylated-STAT-1 levels after 24 h normoxia in control (C) and resistant (R) cells. Protein densities were normalised to β-actin values (mean ± SD, n = 3 independent experiments). **(c)** Expression of genes after exposure to dsRNA synthetic poly I:C (20 μg/mL) for 6 h. **(d)** Expression of genes 48 h after exposure to 0, 6, or 10 Gy radiation. For all qPCR, data represent log-transformed values of fold changes normalised to control cells (measured in triplicates, n = 3 independent experiments, * $P_{adj} < 0.05$, ** $P_{adj} < 0.01$, *** $P_{adj} < 0.001$, 2-way ANOVA with Holm-Sidak correction).

**Supplementary Figure 14.** Resistant cells enhance initial regrowth of mixed xenografts, but not overall survival after single dose radiation. **(a)** Survival curves of PC3 xenografts comprising control (n = 11 mice), 1:1 ctrl:res (n = 10 mice), or resistant cells (n = 11 mice) after 5 Gy irradiation. **(b)** Tumour volumes in initial days of regrowth after radiation (day 0). Data represent mean ± SEM, * $P_{adj} < 0.05$ by 2-way ANOVA with Dunnett's test for multiple comparisons. **(c)** Cell cycle distribution of control and resistant populations isolated from unirradiated PC3 xenografts comprising control (n = 8 mice), mixed (n = 8 mice), or resistant (n = 9 mice) tumours, as measured by flow cytometry. * $P_{adj} < 0.05$, *** $P_{adj} < 0.001$ by 2-way ANOVA with Dunnett's test for multiple comparisons.

SI Fig 1

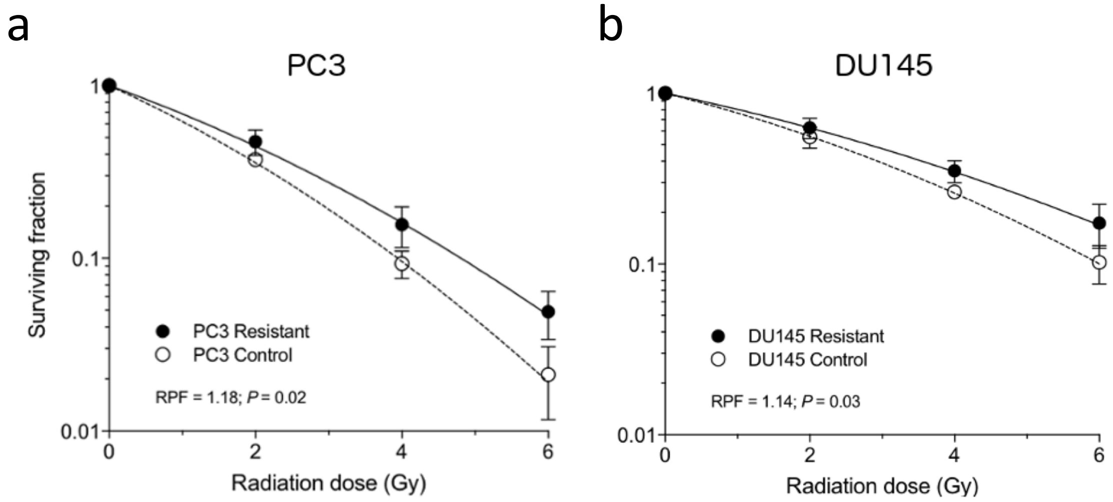
a
PC3
Surviving fraction
1
0.1
0.01
PC3 Resistant
PC3 Control
RPF = 1.18; P = 0.02
0
2
4
6
Radiation dose (Gy)
b
DU145
1
0.1
0.01
DU145 Resistant
DU145 Control
RPF = 1.14; P = 0.03
0
2
4
6
Radiation dose (Gy)

SI Fig 2

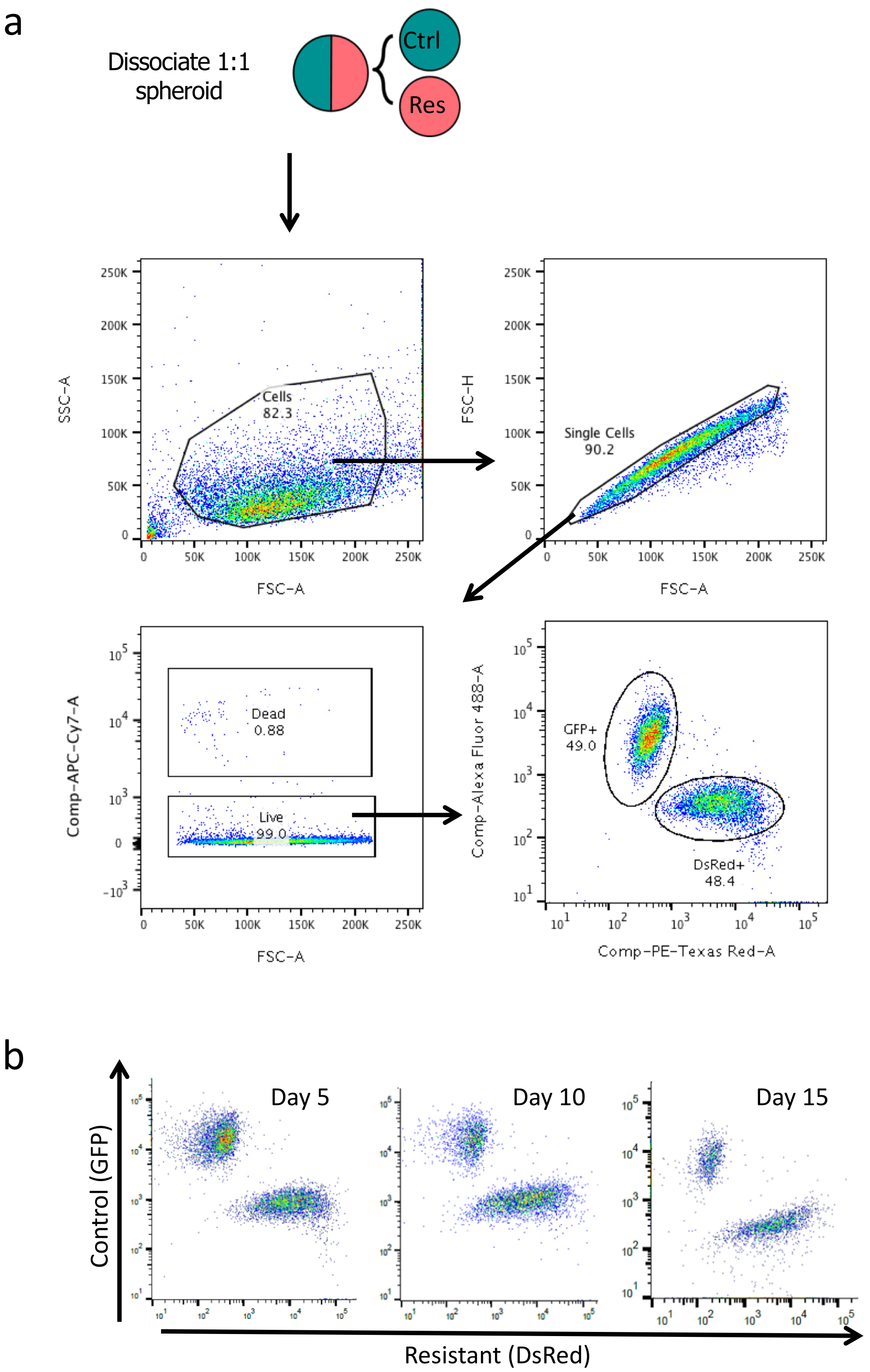
a
Dissociate 1:1 spheroid
Ctrl
Res
Cells 82.3
SSC-A
FSC-A
Single Cells 90.2
FSC-H
Dead 0.88
Live 99.0
Comp-APC-Cy7-A
GFP+ 49.0
DsRed+ 48.4
Comp-Alexa Fluor 488-A
Comp-PE-Texas Red-A
b
Day 5
Day 10
Day 15
Control (GFP)
Resistant (DsRed)

SI Fig 3

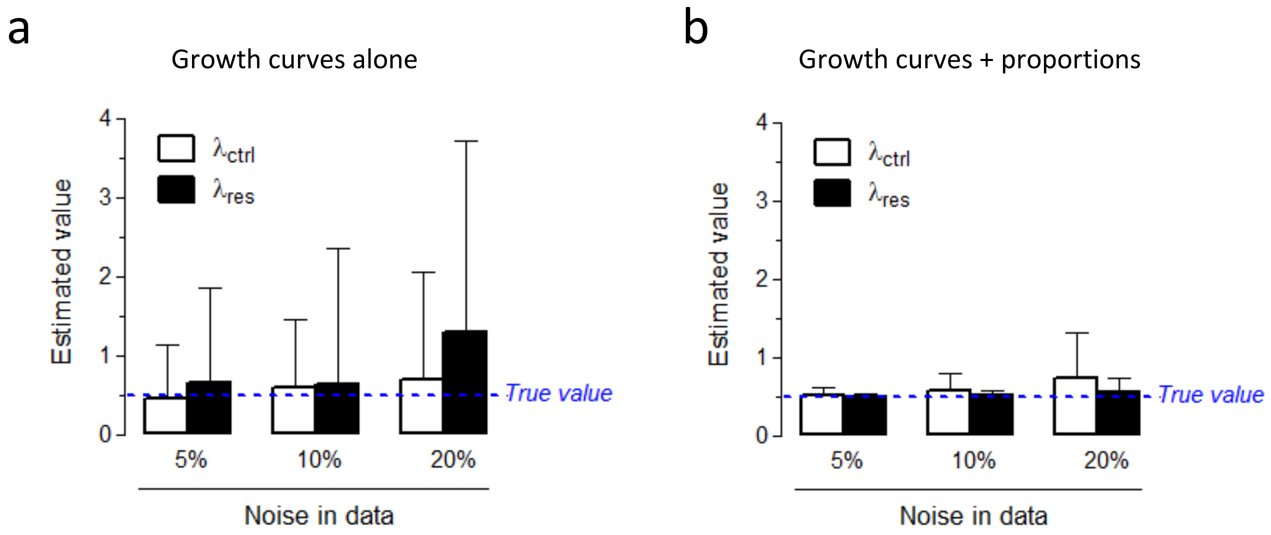
a
Growth curves alone
Estimated value
$\lambda_{ctrl}$
$\lambda_{res}$
True value
5%
10%
20%
Noise in data
b
Growth curves + proportions
Estimated value
$\lambda_{ctrl}$
$\lambda_{res}$
True value
5%
10%
20%
Noise in data

SI Fig 4

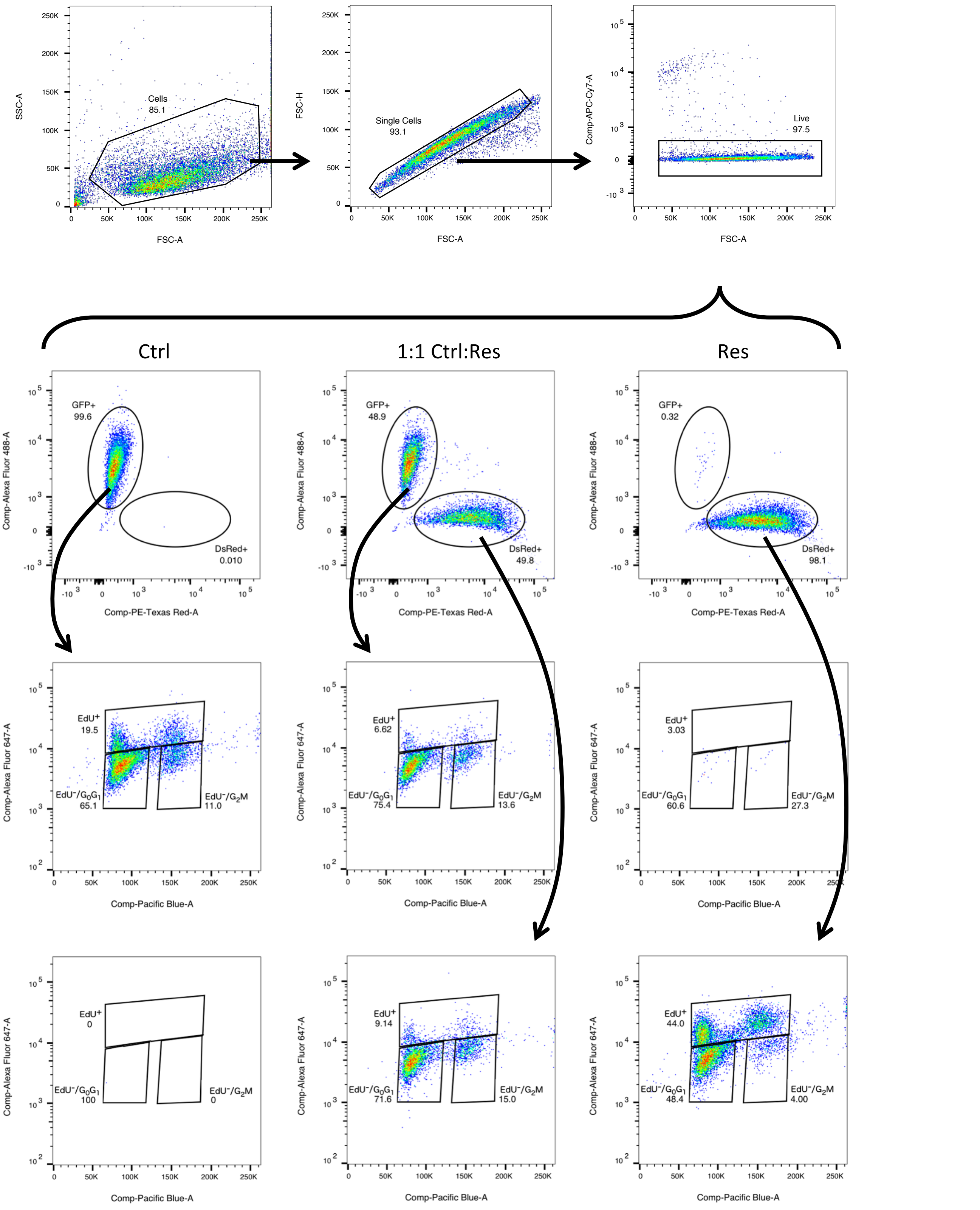

Cells
85.1
Single Cells
93.1
Live
97.5
SSC-A
FSC-A
FSC-H
Comp-APC-Cy7-A
Ctrl
1:1 Ctrl:Res
Res
GFP+
99.6
DsRed+
0.010
GFP+
48.9
DsRed+
49.8
GFP+
0.32
DsRed+
98.1
Comp-Alexa Fluor 488-A
Comp-PE-Texas Red-A
EdU+
19.5
EdU-/G0G1
65.1
EdU-/G2M
11.0
EdU+
6.62
EdU-/G0G1
75.4
EdU-/G2M
13.6
EdU+
3.03
EdU-/G0G1
60.6
EdU-/G2M
27.3
EdU+
0
EdU-/G0G1
100
EdU-/G2M
0
EdU+
9.14
EdU-/G0G1
71.6
EdU-/G2M
15.0
EdU+
44.0
EdU-/G0G1
48.4
EdU-/G2M
4.00
Comp-Alexa Fluor 647-A
Comp-Pacific Blue-A

SI Fig 5

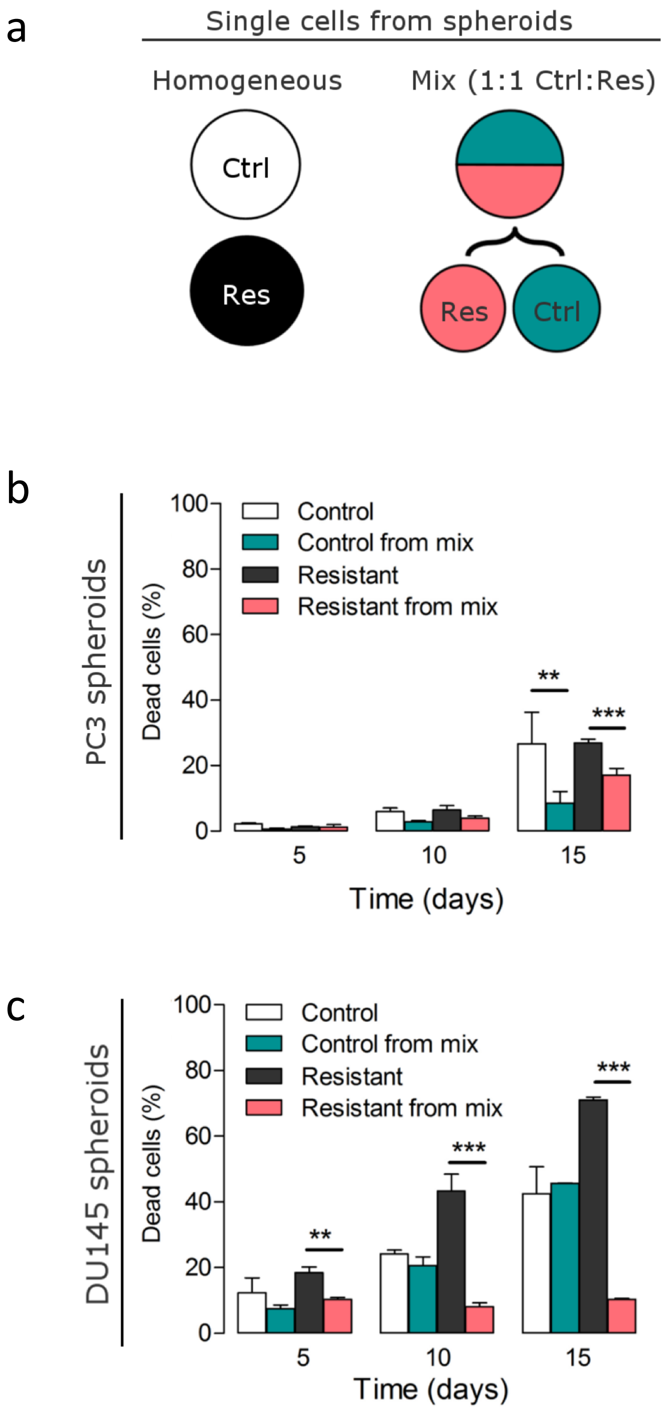

a
Single cells from spheroids
Homogeneous
Mix (1:1 Ctrl:Res)
Ctrl
Res
Res
Ctrl
b
PC3 spheroids
Dead cells (%)
Control
Control from mix
Resistant
Resistant from mix
**
***
Time (days)
c
DU145 spheroids
Dead cells (%)
Control
Control from mix
Resistant
Resistant from mix
**
***
***
Time (days)

SI Fig 6

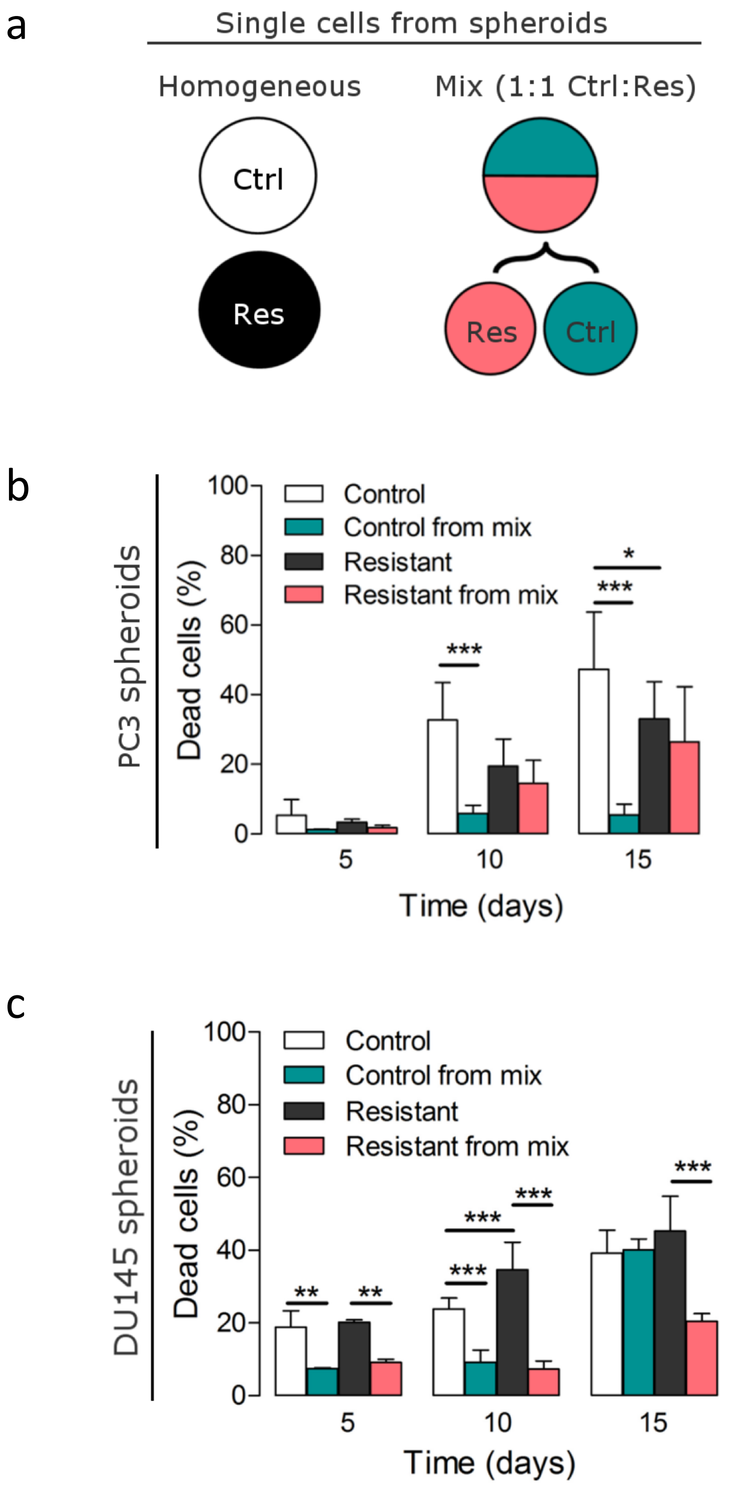

a
Single cells from spheroids
Homogeneous
Mix (1:1 Ctrl:Res)
Ctrl
Res
Res
Ctrl
b
PC3 spheroids
Dead cells (%)
Control
Control from mix
Resistant
Resistant from mix
***
***
*
0
20
40
60
80
100
5
10
15
Time (days)
c
DU145 spheroids
Dead cells (%)
Control
Control from mix
Resistant
Resistant from mix
**
**
***
***
***
***
0
20
40
60
80
100
5
10
15
Time (days)

### SI Fig 7

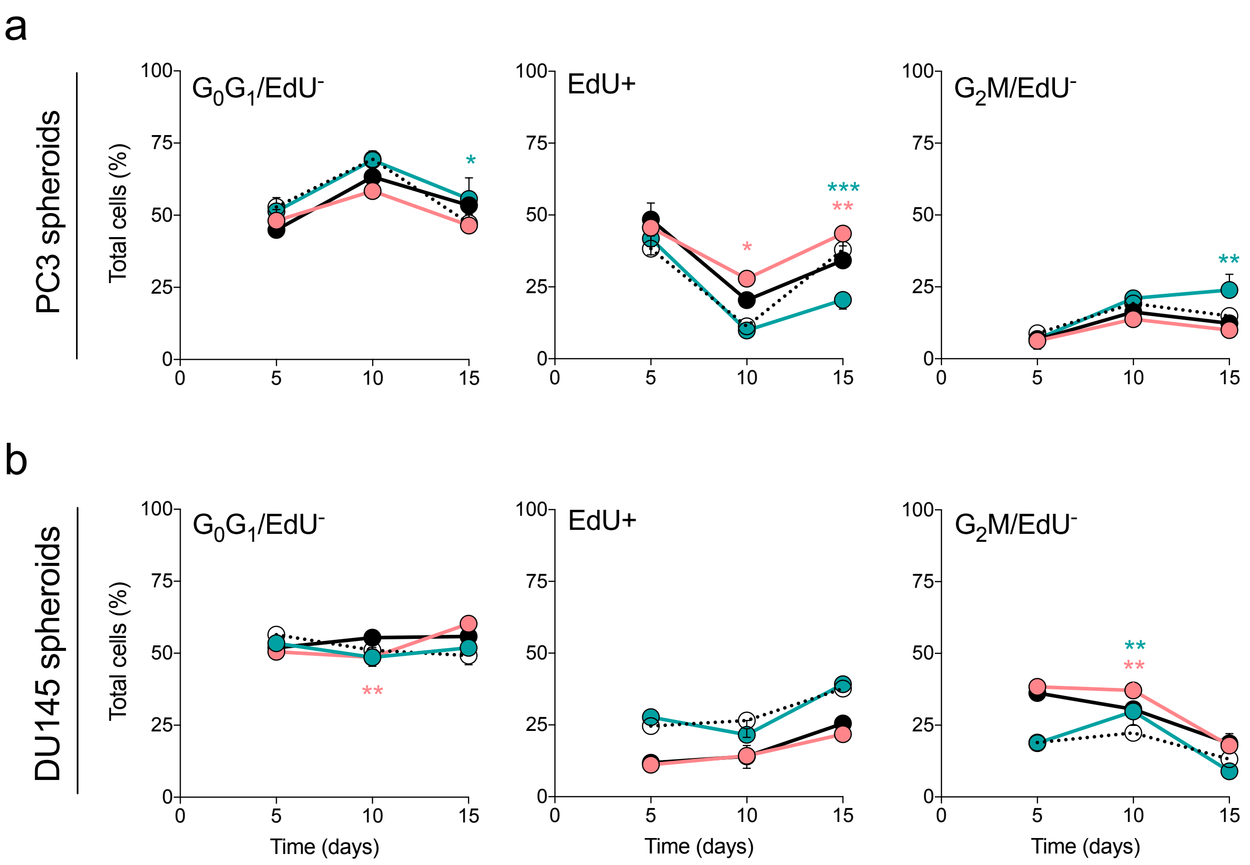

a
PC3 spheroids
Total cells (%)
$G_0G_1$/EdU$^-$
EdU+
$G_2$M/EdU$^-$
b
DU145 spheroids
Total cells (%)
$G_0G_1$/EdU$^-$
EdU+
$G_2$M/EdU$^-$
Time (days)
Time (days)
Time (days)

SI Fig 8

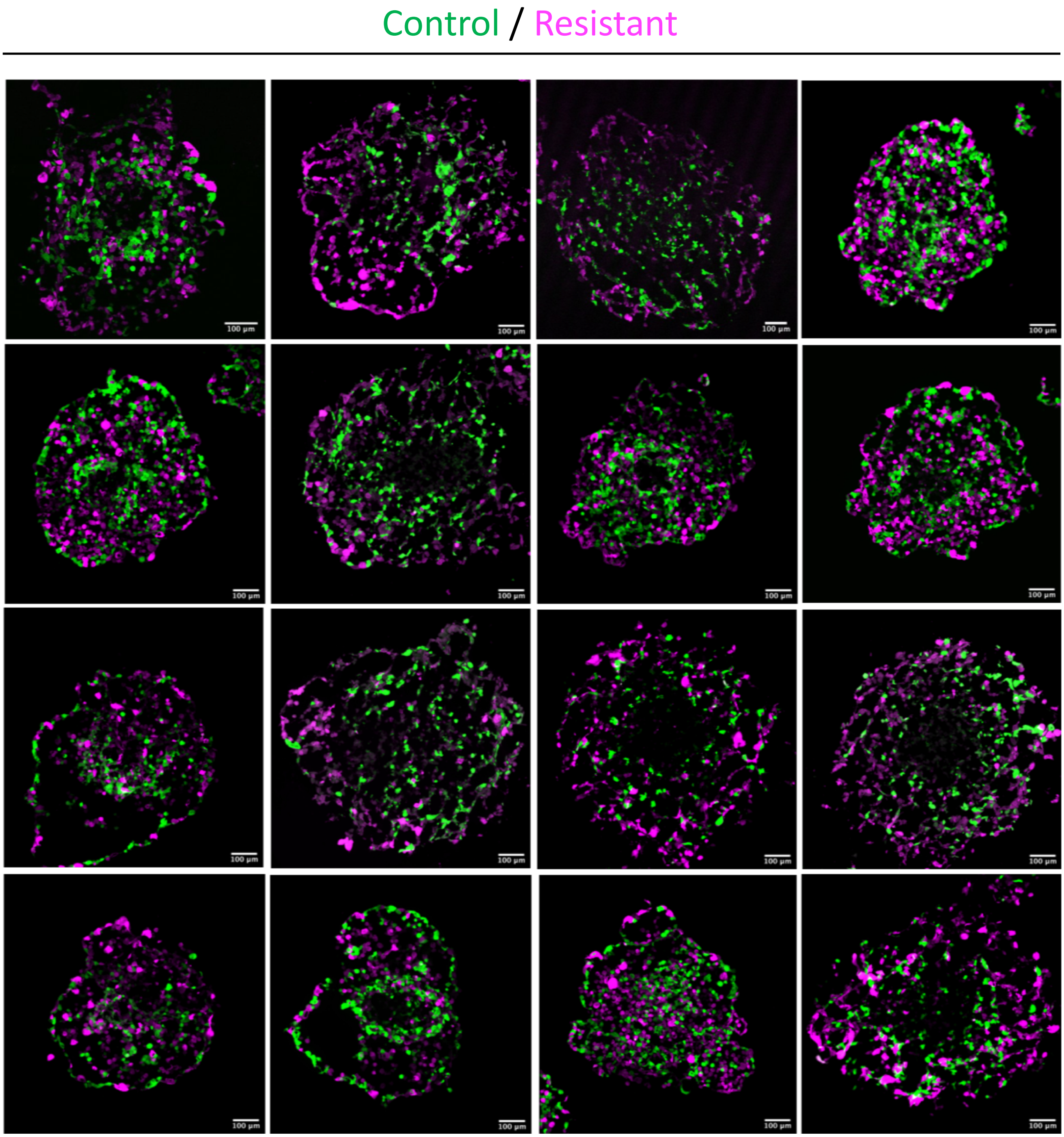

### SI Fig 9

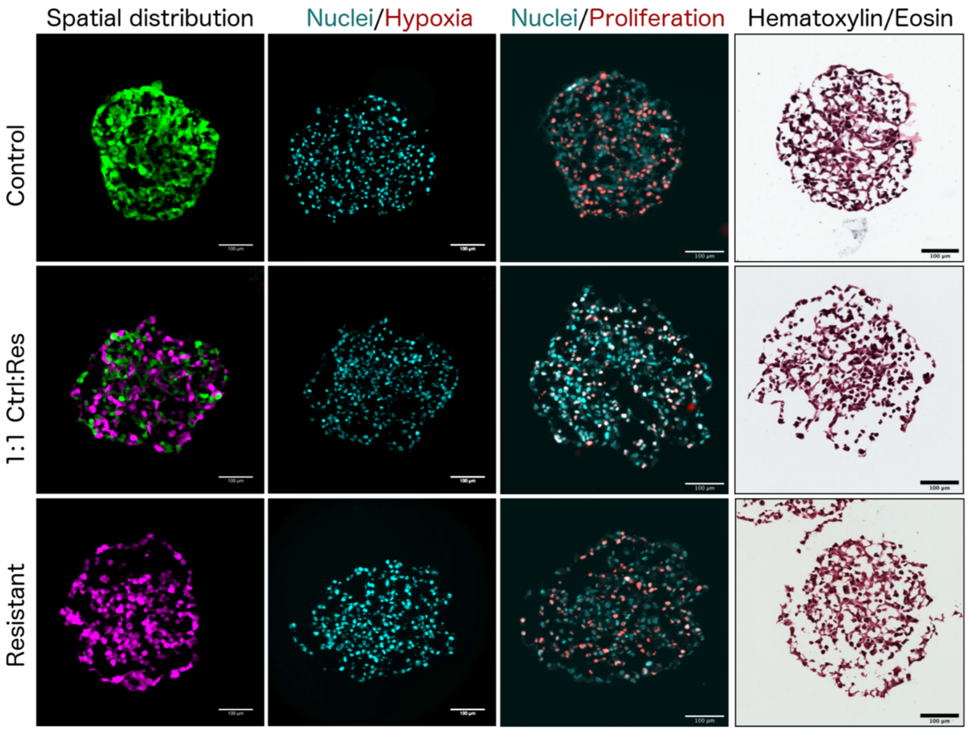
Spatial distribution
Nuclei/Hypoxia
Nuclei/Proliferation
Hematoxylin/Eosin
Control
1:1 Ctrl:Res
Resistant

SI Fig 10

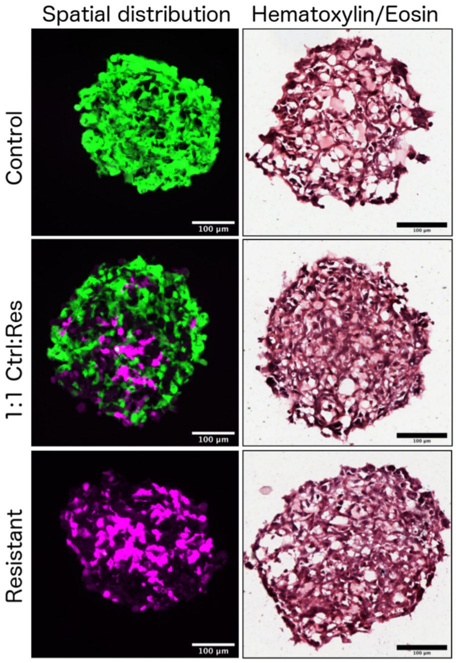

Spatial distribution
Hematoxylin/Eosin
Control
1:1 Ctrl:Res
Resistant
100 µm
100 µm
100 µm

SI Fig 11

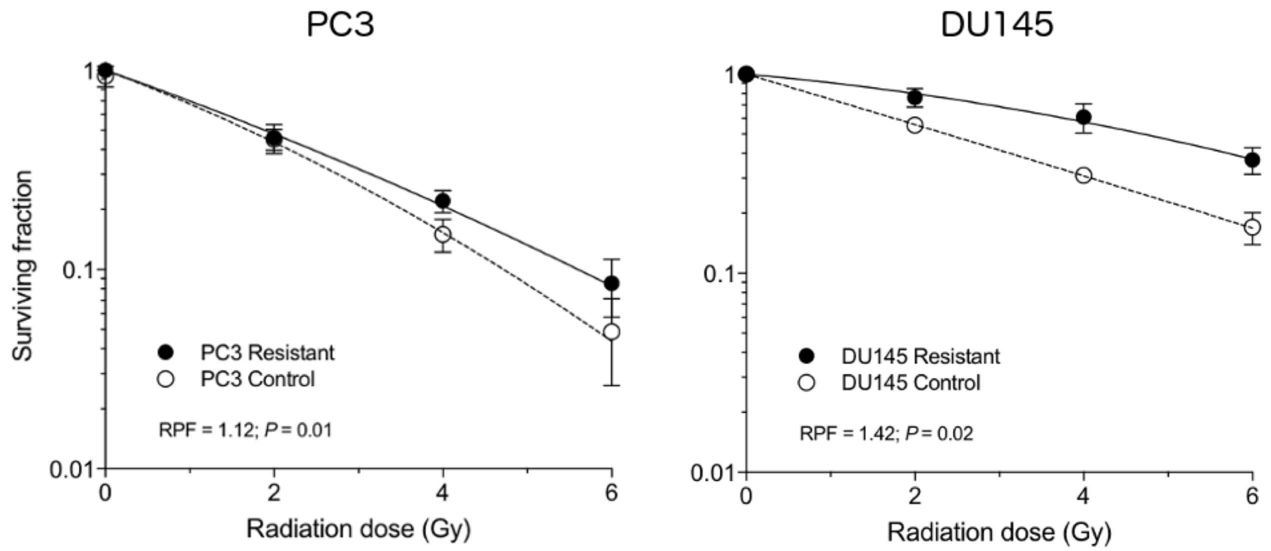

PC3
DU145
Surviving fraction
1
0.1
0.01
0
2
4
6
PC3 Resistant
PC3 Control
RPF = 1.12; P = 0.01
DU145 Resistant
DU145 Control
RPF = 1.42; P = 0.02
Radiation dose (Gy)

SI Fig 12

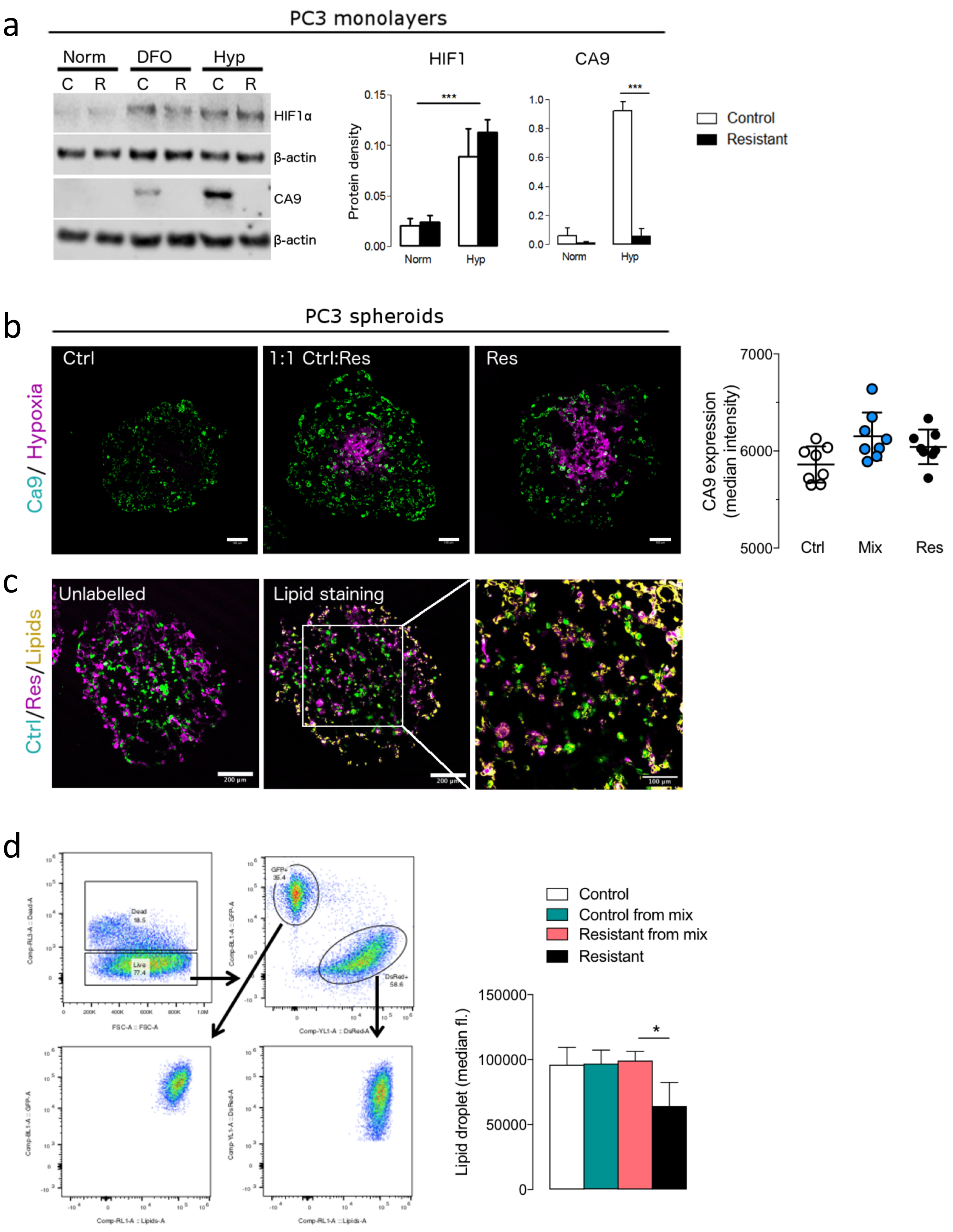

a
PC3 monolayers
Norm
DFO
Hyp
C R C R C R
HIF1α
β-actin
CA9
β-actin
HIF1
CA9
Protein density
Norm
Hyp
Control
Resistant
b
PC3 spheroids
Ca9/ Hypoxia
Ctrl
1:1 Ctrl:Res
Res
CA9 expression (median intensity)
Ctrl
Mix
Res
c
Ctrl/Res/Lipids
Unlabelled
Lipid staining
200 µm
100 µm
d
Control
Control from mix
Resistant from mix
Resistant
Lipid droplet (median fl.)

SI Fig 13

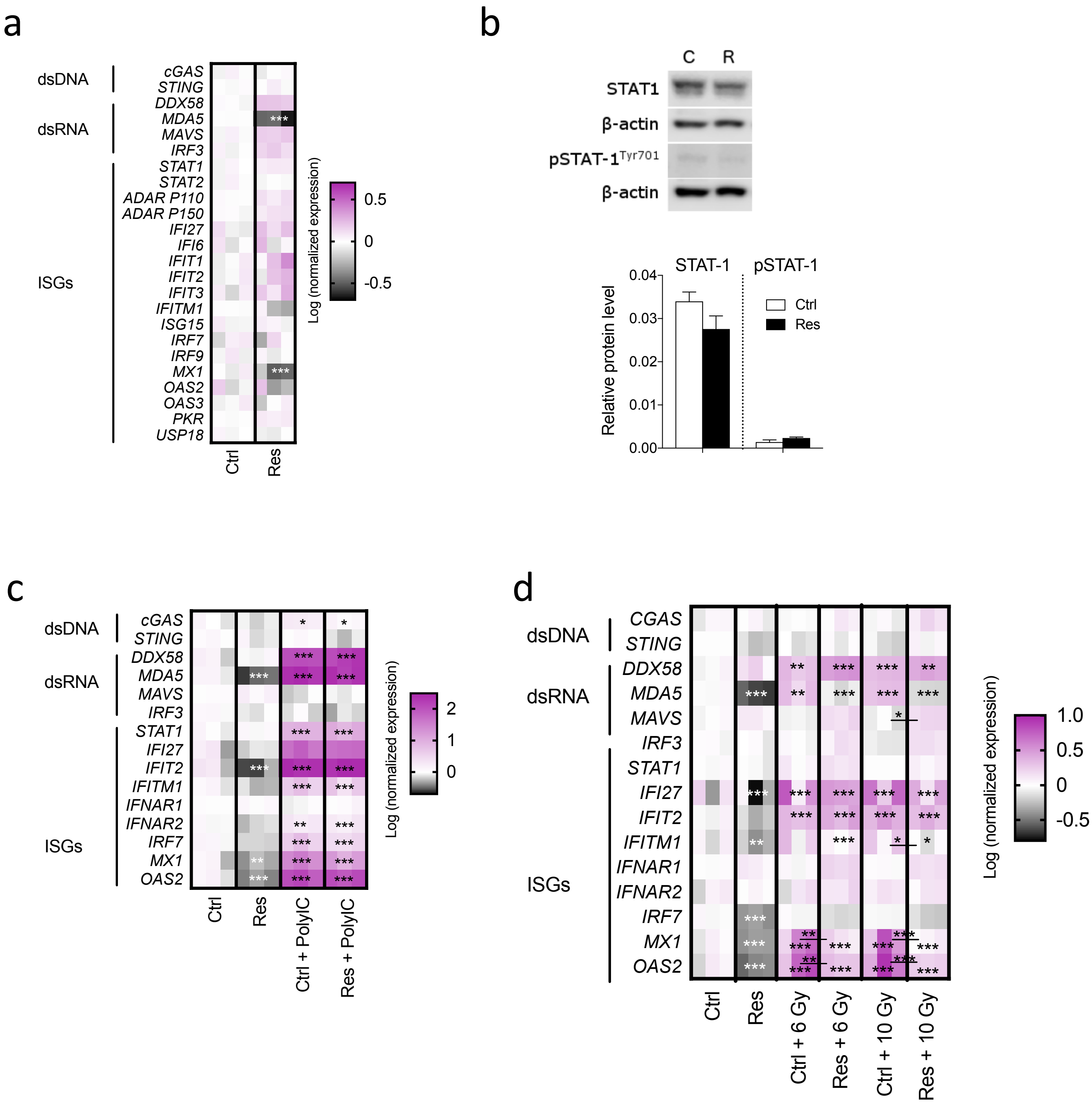
a
dsDNA
dsRNA
ISGs
cGAS
STING
DDX58
MDA5
MAVS
IRF3
STAT1
STAT2
ADAR P110
ADAR P150
IFI27
IFI6
IFIT1
IFIT2
IFIT3
IFITM1
ISG15
IRF7
IRF9
MX1
OAS2
OAS3
PKR
USP18
Ctrl
Res
Log (normalized expression)
0.5
0
-0.5
b
C
R
STAT1
β-actin
pSTAT-1Tyr701
β-actin
STAT-1
pSTAT-1
Relative protein level
0.04
0.03
0.02
0.01
0.00
Ctrl
Res
c
dsDNA
dsRNA
ISGs
cGAS
STING
DDX58
MDA5
MAVS
IRF3
STAT1
IFI27
IFIT2
IFITM1
IFNAR1
IFNAR2
IRF7
MX1
OAS2
Ctrl
Res
Ctrl + PolyIC
Res + PolyIC
Log (normalized expression)
2
1
0
d
dsDNA
dsRNA
ISGs
CGAS
STING
DDX58
MDA5
MAVS
IRF3
STAT1
IFI27
IFIT2
IFITM1
IFNAR1
IFNAR2
IRF7
MX1
OAS2
Ctrl
Res
Ctrl + 6 Gy
Res + 6 Gy
Ctrl + 10 Gy
Res + 10 Gy
Log (normalized expression)
1.0
0.5
0
-0.5

SI Fig 14

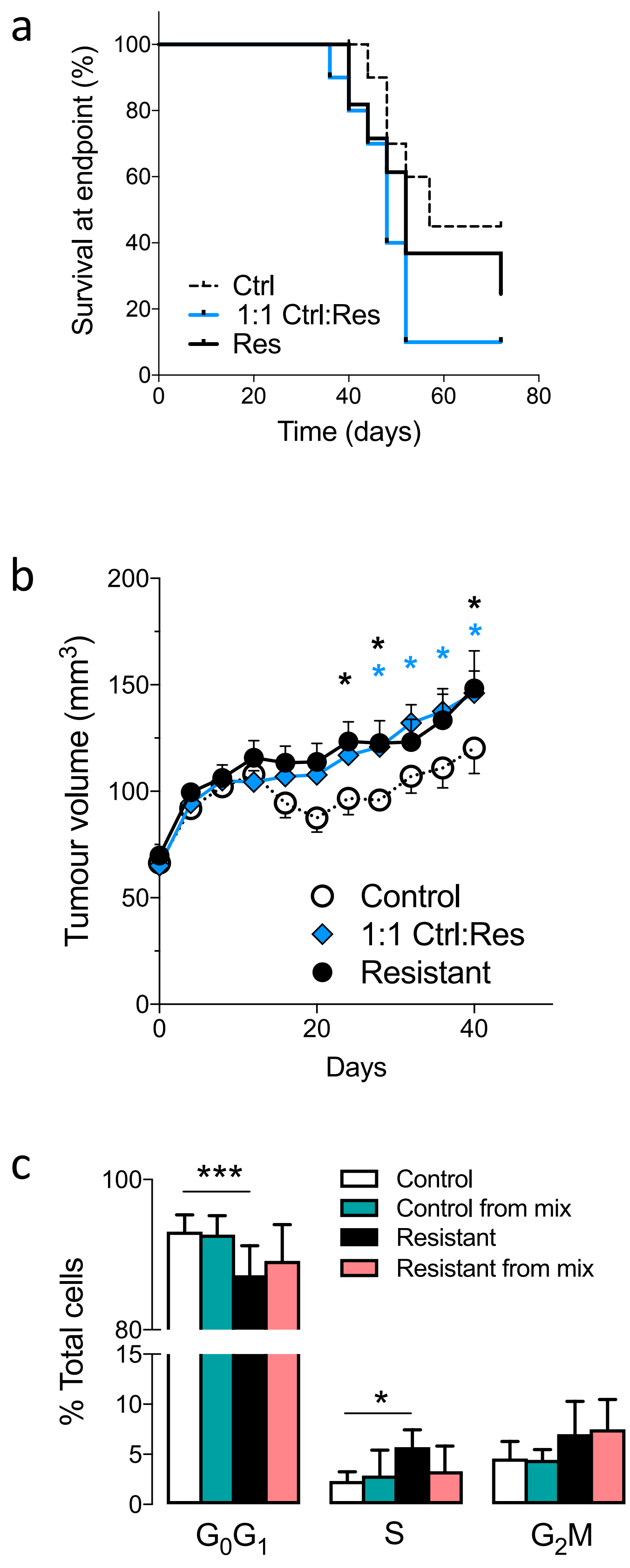
a
Survival at endpoint (%)
100
80
60
40
20
0
0
20
40
60
80
Time (days)
Ctrl
1:1 Ctrl:Res
Res
b
Tumour volume (mm3)
200
150
100
50
0
0
20
40
Days
Control
1:1 Ctrl:Res
Resistant
c
% Total cells
100
80
15
10
5
0
***
*
Control
Control from mix
Resistant
Resistant from mix
G0G1
S
G2M

### Supplementary Methods

#### 1 Spatially-averaged mathematical models

##### 1.1 Logistic model of homogeneous spheroid growth

The growth of tumour spheroids typically begins with a phase of exponential growth which is followed by an intermediate linear growth phase and eventually superseded by a slower approach to an equilibrium size at which the net rates of cell death and proliferation across the tumour volume are balanced. Growth laws that have been proposed to describe this process include the Gompertz, Bertalanffy and logistic models [5]. Here we use the latter to model the growth of homogeneous spheroids. The logistic growth law states that the rate of change of the tumour spheroid volume $V(t)$ (mm$^3$) at time $t$ (day) is given by

$$\frac{dV}{dt} = rV\left(1 - \frac{V}{K}\right), \text{ with } V(t=0) = V_0, \tag{1}$$

where the parameters $r > 0$ (day$^{-1}$) and $K > 0$ (mm$^3$) represent the growth rate and carrying capacity, or equilibrium size, of the tumour and $V_0 > 0$ (mm$^3$) denotes the spheroid volume at $t = 0$. The initial value problem (1) can be solved to give

$$V(t) = \frac{V_0 K e^{rt}}{K + V_0(e^{rt} - 1)}.$$

##### 1.2 Lotka-Volterra model of heterogeneous spheroid growth

We assume that when the control and resistant cell populations are co-cultured to form heterogeneous spheroids, their growth dynamics and interactions can be viewed as a Lotka-Volterra system. If we denote by $V_C(t)$ and $V_R(t)$ the volumes of the tumour occupied by the control and resistant cells respectively at time $t$, then their evolution is described by the following system of ordinary differential equations:

$$\begin{aligned} \frac{dV_C}{dt} &= V_C \left[ r_C \left( 1 - \frac{V_C}{K_C} \right) - \lambda_R V_R \right], \\ \frac{dV_R}{dt} &= V_R \left[ r_R \left( 1 - \frac{V_R}{K_R} \right) - \lambda_C V_C \right]. \end{aligned} \tag{2}$$

In Eqs. (2), the positive parameters $r_C$ (day$^{-1}$) and $r_R$ (day$^{-1}$) are the intrinsic growth rates of the two populations and the positive parameters $K_C$ (mm$^3$) and $K_R$ (mm$^3$) their carrying capacities (i.e., the equilibrium volumes to which the control and resistant populations would evolve if they were cultured as homogeneous spheroids). The parameters $\lambda_C$ (day$^{-1}$ mm$^{-1}$) and $\lambda_R$ (day$^{-1}$ mm$^{-1}$) respectively describe the effect that the control cells have on the resistant cells, and vice versa. In the absence of prior knowledge about the nature of the interactions between the two cell populations, we do not restrict $\lambda_C$ and $\lambda_R$ to be positive or negative. Indeed, as the signs of $\lambda_C$ and $\lambda_R$ vary we can distinguish six types of interactions: these are summarised in Table 1. We note that, if $\lambda_C = \lambda_R = 0$, then Eqs. (2) reduce to two logistic equations for $V_C$ and $V_R$ similar to Eq. (1).

**Table 1:** Potential interactions between tumour cell populations. The table summarises the biological effects that the control and resistant cell populations can have on each other, together with the corresponding signs of the parameters $\lambda_C$ and $\lambda_R$ in Eqs. (2).

| **Type of interaction** | **Effect of control cells on resistant cells** | **Effect of resistant cells on control cells** | **Sign of** $\lambda_C$ | **Sign of** $\lambda_R$ |
|---|---|---|---|---|
| ***Competition*** | Detrimental | Detrimental | $\lambda_C > 0$ | $\lambda_R > 0$ |
| ***Amensalism*** | Detrimental | No effect | $\lambda_C > 0$ | $\lambda_R = 0$ |
| ***Antagonism*** | Detrimental | Beneficial | $\lambda_C > 0$ | $\lambda_R < 0$ |
| ***Neutralism*** | No effect | No effect | $\lambda_C = 0$ | $\lambda_R = 0$ |
| ***Commensalism*** | No effect | Beneficial | $\lambda_C = 0$ | $\lambda_R < 0$ |
| ***Mutualism*** | Beneficial | Beneficial | $\lambda_C < 0$ | $\lambda_R < 0$ |

In order to arrive at a well defined initial value problem, Eqs. (2) are supplemented by the following initial conditions:

$$V_C(t=0) = V_{C0} \qquad \text{and} \qquad V_R(t=0) = V_{R0}. \tag{3}$$

For a homogeneous spheroid, whose growth can be modelled using Eq. (1), there is an obvious physical interpretation of the carrying capacity parameter $K$ – it is the maximum volume of the spheroid that can be supported by its environment. For a heterogeneous spheroid, such as that modelled with Eqs. (2), the spheroid's saturation size cannot be described by a single parameter. Indeed the different interactions that may exist between co-cultured cell populations manifest themselves in a multitude of possible growth regimes. Nonetheless, we can consider some simple cases. Let us first denote by $K_T$ the carrying capacity of a heterogeneous spheroid, when it exists. If $\lambda_C = \lambda_R = 0$ then it is straightforward to show that $K_T = K_C + K_R$. In the case of competition, when $\lambda_C > 0$ and $\lambda_R > 0$, we would expect $K_T < K_C + K_R$. Similarly, when the populations support each other, that is when $\lambda_C < 0$ and $\lambda_R < 0$ (mutualism), we anticipate that $K_T > K_C + K_R$.

For logistic growth, the carrying capacity $K$ corresponds to a stable steady state (see Eq. (1)). Similarly, the co-existence equilibrium solutions of Eqs. (2), where they exist and are stable, define the saturation sizes of heterogeneous spheroids. By setting time-derivatives equal to zero in Eqs. (2), it is straightforward to show that Eqs. (2) possess up to four steady states:

1. $(V_C, V_R) = (0, 0)$: tumour elimination;
2. $(V_C, V_R) = (K_C, 0)$: homogeneous tumour spheroid, with radio-resistant cells eliminated;
3. $(V_C, V_R) = (0, K_R)$: homogeneous tumour spheroid, with radio-sensitive cells eliminated;
4. $(V_C, V_R) = (V_C^*, V_R^*)$: coexistence of both cell populations, where

$$V_C^* = \frac{\left(\frac{1}{K_R} - \frac{\lambda_R}{r_C}\right)}{\left(\frac{1}{K_R K_C} - \frac{\lambda_R \lambda_C}{r_R r_C}\right)} \quad \text{and} \quad V_R^* = \frac{\left(\frac{1}{K_C} - \frac{\lambda_C}{r_R}\right)}{\left(\frac{1}{K_R K_C} - \frac{\lambda_R \lambda_C}{r_R r_C}\right)},$$

$$\Rightarrow V_T = V_C^* + V_R^* = \frac{\left(\frac{1}{K_R} + \frac{1}{K_C}\right) - \left(\frac{\lambda_R}{r_C} + \frac{\lambda_C}{r_R}\right)}{\left(\frac{1}{K_R K_C} - \frac{\lambda_R \lambda_C}{r_R r_C}\right)}.$$

Steady states 1, 2 and 3 exist for all choices of the interaction parameters $\lambda_C$ and $\lambda_R$. By contract, the coexistence steady state is only physically realistic and stable for certain combinations of values of $r_C$, $r_R$, $K_C$, $K_R$, $\lambda_C$ and $\lambda_R$. In particular, if we define $\eta_R = \frac{\lambda_R K_R}{r_C}$ and $\eta_C = \frac{\lambda_C K_C}{r_R}$ then the coexistence steady state exists in the shaded regions of Fig. 1 where

$$V_C^* = \frac{(1 - \eta_R)}{(1 - \eta_R \eta_C)} K_C \quad \text{and} \quad V_R^* = \frac{(1 - \eta_C)}{(1 - \eta_R \eta_C)} K_R,$$

$$\Rightarrow V_T = \frac{(1 - \eta_R) K_C + (1 - \eta_C) K_R}{(1 - \eta_R \eta_C)}.$$

We note also that if $\eta_R = \eta_C = \eta$, say, then $V_T = (1 + \eta)(K_C + K_R)$, in which case $0 < V_T < (K_C + K_R)$ for $-1 < \eta < 0$ and $V_T > (K_C + K_R)$ for $0 < \eta$.

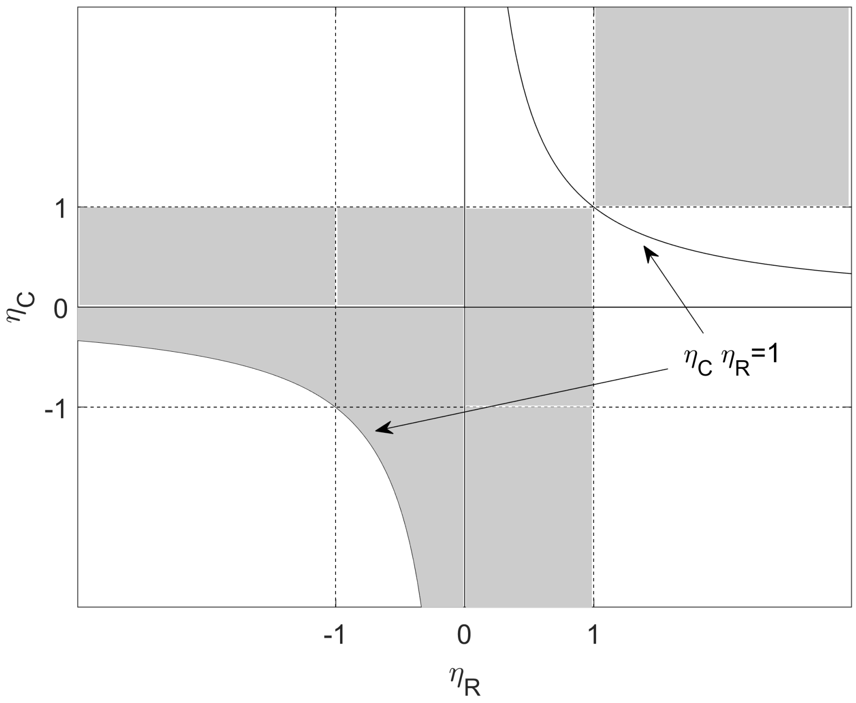


**Figure 1:** The coexistence steady state is physically realistic and stable in the shaded regions of the $\eta_R\eta_C$-plane, where $\eta_R = \frac{\lambda_R K_R}{r_C}$ and $\eta_C = \frac{\lambda_C K_C}{r_R}$.

#### 1.3 Parameter estimation

##### 1.3.1 Logistic model

The values of parameters $\theta_{hom} = (r, K, V_0)$ in Eq. (1) were estimated by minimising the weighted sum of squared residuals between experimental measurements of spheroid volume and the solution to Eq. (1) for given values of $\theta_{hom}$. In mathematical terms, we sought $\hat{\theta}_{hom}$ such that

$$\hat{\theta}_{hom} = \arg\min_{\theta_{hom}} \sum_{i=1}^{N} \left[ \frac{V_i^{data} - V^{model}(t_i; \theta_{hom})}{V^{model}(t_i; \theta_{hom})} \right]^2 \tag{4}$$

where $V_i^{data}$ are the volume measurements taken at times $t_i$ ($i = 6$ for PC3 and $i = 9$ for DU145 spheroids, see Fig. 1b in the main text) and $V^{model}(t_i; \theta_{hom})$ are model outputs at $t_i$ for given values of the model parameters $\theta_{hom}$.

We remark that there is no consensus regarding how to weight the data when minimising the sum of squared residuals. In equation (4), we follow [1] and use the model to weight the data.

##### 1.3.2 Lotka-Volterra model

The values of the parameters $r_C, r_R, K_C, K_R, V_{C0}$ and $V_{R0}$ describing the growth rates, carrying capacities and initial volumes for each cell population were estimated using the volume data collected from the homogeneous spheroids, and were subsequently fixed in Eqs. (2). Thus the only unknown parameters in Eqs. (2) were the interaction parameters $\theta_{het} = (\lambda_C, \lambda_R)$. These were estimated by seeking $\theta_{het}$ that minimises the following sum of squared residuals:

$$\sum_{j=1}^{M} \left[ \frac{V_C^{data}(t_j) - V_C^{model}(t_j;\theta_{het})}{V_C^{model}(t_j;\theta_{het})} \right]^2 + \sum_{j=1}^{M} \left[ \frac{V_R^{data}(t_j) - V_R^{model}(t_j;\theta_{het})}{V_R^{model}(t_j;\theta_{het})} \right]^2 + \sum_{i=1}^{N} \left[ \frac{V_T^{data}(t_i) - V_T^{model}(t_i;\theta_{het})}{V_T^{model}(t_i;\theta_{het})} \right]^2, \tag{5}$$

where $V_C^{data}(t_j)$ and $V_R^{data}(t_j)$ respectively represent the volumes of the control and resistant populations within the heterogeneous spheroids at times $t_j$, $V_C^{model}(t_j;\theta_{het})$ and $V_R^{model}(t_j;\theta_{het})$ are solutions to Eqs. (2) at $t_j$ for given values of $\theta_{het}$, $V_T^{data}(t_i) = V_C^{data}(t_i) + V_R^{data}(t_i)$ are the total volumes of the heterogeneous spheroids at times $t_i$ and $V_T^{model}(t_i;\theta_{het}) = V_C^{model}(t_i;\theta_{het}) + V_R^{model}(t_i;\theta_{het})$ are the corresponding solutions to Eqs. (2) at $t_i$ for $\theta_{het}$. The times $t_i$ at which total spheroid volumes were measured are shown in Fig. 1b (main text). The times $t_j$ at which the proportions of the control and resistant cells were measured are $t_j = (5, 10, 15, 19)$ (days) for PC3 and $t_j = (5, 10, 15)$ (days) for DU145 spheroids.

The minimisation problems (4) and (5) were implemented and solved in MATLAB using a non-linear least squares solver *lsqnonlin* which is an implementation of the trust-region-reflective iterative optimisation algorithm [3]. Since *lsqnonlin* is a local solver we use it in combination with *MultiStart*, a function that runs the local solver from a number of randomly selected starting points within a prescribed region of parameter space, thus ensuring more thorough exploration of the parameter space.

#### 1.4 Radiation response modelling for theoretical study

To simulate growth of heterogeneous spheroids after exposure to radiation we used Eqs. (2) for a range of values of $\lambda_C$ and $\lambda_R$ (with other parameter values fixed) to grow *in silico* spheroids until they reached a volume of $V_{IR} = 0.9\text{mm}^3$. We then simulated radiation damage by calculating the surviving fraction $\sigma$ of each population according to the linear-quadratic model [9]. In particular we set

$$V_\gamma(t_{IR^+}) = \sigma_\gamma V_\gamma(t_{IR^-}) \tag{6}$$

where

$$\sigma_\gamma = e^{-(\alpha_\gamma d + \beta_\gamma d^2)} \tag{7}$$

for $\gamma \in \{C, R\}$. In Eqs. (6) and (7) $t_{IR^{\pm}}$ represent the times just before ($t_{IR^{+}}$) and just after ($t_{IR^{-}}$) irradiation, and $\alpha_\gamma$ (Gy$^{-1}$) and $\beta_\gamma$ (Gy$^{-2}$) are the lethal lesions made by one- and two-track actions of radiation dose $d$. Following radiation we used Eqs. (2) to regrow the *in silico* spheroids until they reached their pre-irradiation volume of 0.9mm$^3$. Values of the parameters $\alpha_\gamma$ and $\beta_\gamma$ were estimated for each cell population using data collected from the clonogenic assays. The resulting growth curves were used to generate surface plots representing the regrowth time (Figure 4 in the main text).

### 2 Spatially resolved computational model

We developed a 2D hybrid cellular automaton model of avascular tumor growth and assumed that it is representative of the changes in the size and structure of a 2D cross-section through a 3D tumor spheroid suspended in culture medium. The hybrid CA model couples a set of automaton elements, each with size $l \times l$, arranged on a regular $L \times L$ grid to a reaction-diffusion equation (RDE) describing the spatial distribution of a growth-rate-limiting nutrient which is supplied from the culture medium surrounding the spheroid. Unless otherwise stated, we fix $L = 200$ and take the size of each automaton element to represent the size of an average PC3 cell ($l = 18$ µm). Then the grid represents a region of size $0.36 \times 0.36$ cm$^2$. We consider oxygen ($O_2$) to be the growth-rate-limiting nutrient and model its concentration explicitly. The reaction-diffusion equation is discretised and solved on the same 2D grid as the cellular automaton (see Section 2.1 for details). Each automaton element is occupied either by a cell or by culture medium and has associated with it an oxygen concentration. The behaviour of cells on the grid depends on their local $O_2$ concentration and the occupancy of their neighbourhood. These factors determine the rate at which cells consume $O_2$, and whether they proliferate, become quiescent or die.

In the remainder of this section we describe how the reaction-diffusion equation was discretised and solved, outline the rules governing our CA model and summarise the parameters (and parameter values) used in the model simulations.

#### 2.1 Oxygen dynamics

A reaction-diffusion equation models the concentration of oxygen within our simulation domain. If we denote the oxygen concentration at location $\mathbf{x}$ and time $t$ by $c(\mathbf{x}, t)$ (mol cm$^{-3}$) then its evolution is described by

$$\frac{\partial c(\mathbf{x}, t)}{\partial t} = D\nabla^2 c(\mathbf{x}, t) - \Gamma(\mathbf{x}, t), \tag{8}$$

where $D$ is the (constant) oxygen diffusion coefficient (cm$^2$ s$^{-1}$) and $\Gamma(\mathbf{x}, t)$ is the oxygen consumption rate (mol cm$^{-3}$ s$^{-1}$). In what follows, we fix

$$\Gamma(\mathbf{x}, t) = \begin{cases} \kappa_{\mathcal{P}_\gamma} & \text{if } \mathbf{x} \text{ is occupied by a proliferating cell from population } \gamma, \\ \kappa_{\mathcal{Q}_\gamma} & \text{if } \mathbf{x} \text{ is occupied by a quiescent cell from population } \gamma, \\ 0 & \text{otherwise,} \end{cases} \tag{9}$$

where $0 < \kappa_{\mathcal{Q}} < \kappa_{\mathcal{P}}$ denote respectively the constant rates at which quiescent and proliferating cells consume oxygen (units: moles cm$^{-3}$ s$^{-1}$). Equation (8) is supplemented by the following initial and boundary conditions

$$c(x, y, 0) = c_\infty, \tag{10}$$

$$c(0, y, t) = c(L, y, t) = c(x, 0, t) = c(x, L, t) = c_\infty \tag{11}$$

where $L$ is the domain length and $c_\infty$ is the background $O_2$ concentration. Equations (8)–(11) describe a situation in which $O_2$ diffuses from the boundaries of a (rectangular) Petri dish (where it is maintained at fixed levels) into the culture medium where it is consumed by tumour cells. Although in theory the $O_2$ concentration could become negative, in practice this does not happen because cells residing in low, but nonzero, $O_2$ conditions become necrotic and do not consume $O_2$.

Eqs. (8)–(11) are nondimensionalised, discretised using central differences for the spatial derivatives and forward differences for the time derivatives, and then solved using an explicit Euler scheme in MATLAB.

#### 2.2 Cellular automaton rules

##### 2.2.1 Cell status

Each automaton element at location $\mathbf{x} = (x, y)$ and time $t$ can be considered a dynamical variable with a *state* and *neighbourhood.* Possible *states* include proliferating ($\mathcal{P}_\gamma$), quiescent ($\mathcal{Q}_\gamma$), necrotic ($\mathcal{N}_\gamma$) and empty ($\mathcal{E}$) cells (empty automaton elements are assumed to contain culture medium) which enables us to represent commonly observed features of tumour spheroids which include the formation of quiescent/hypoxic and necrotic regions in response to oxygen levels. We define quiescent cells to be viable cells that are not actively progressing through the cell cycle. These cells are in the $G_0$ phase and awaiting restoration of favourable conditions so that they can re-enter the cell cycle. This CA can be extended to model the growth of heterogeneous spheroids by introducing more states. For example, by defining $\gamma \in (\mathcal{C}, \mathcal{R})$ *states* becomes $\{\mathcal{P}_\mathcal{C}, \mathcal{P}_\mathcal{R}, \mathcal{Q}_\mathcal{C}, \mathcal{Q}_\mathcal{R}, \mathcal{N}_\mathcal{C}, \mathcal{N}_\mathcal{R}, \mathcal{E}\}$ allowing us to model the growth of heterogeneous spheroids consisting of control ($\mathcal{C}$) and resistant ($\mathcal{R}$) cells.

The *state* of a cell at location $\mathbf{x}$ and time $t$, denoted as $state(\mathbf{x}, t)$, is determined by the concentration of oxygen, $c(\mathbf{x}, t)$, at that location. If we define by $c_{\mathcal{Q}\gamma}$ and $c_{\mathcal{N}\gamma}$ the threshold oxygen levels below which cells of type $\gamma$ become quiescent and necrotic, respectively, then we have:

- if $c_\infty \geq c(\mathbf{x}, t) > c_{\mathcal{Q}\gamma}$ then $state(\mathbf{x}, t) = \mathcal{P}_\gamma$,
- if $c_{\mathcal{Q}_\gamma} \geq c(\mathbf{x}, t) > c_{\mathcal{N}_\gamma}$ then $state(\mathbf{x}, t) = \mathcal{Q}_\gamma$,
- if $c_{\mathcal{N}_\gamma} \geq c(\mathbf{x}, t) \geq 0$ then $state(\mathbf{x}, t) = \mathcal{N}_\gamma$.

In addition to a *state*, every automaton element has a *neighbourhood* associated with it. The *neighbourhood* provides local information about the cell's surroundings and can impact the cell's next *state*. The two most commonly used neighbourhoods in CA models are the von Neumann and Moore neighbourhoods (Fig. 2). The number and location of neighbours can have a significant impact on cell-cell communication. Since it has been previously demonstrated that the Moore neighbourhood can minimise artefacts associated with lattice anisotropies [10] we use the first order Moore neighbourhood. Thus, for a 2D model, every cell communicates with its eight nearest neighbours.

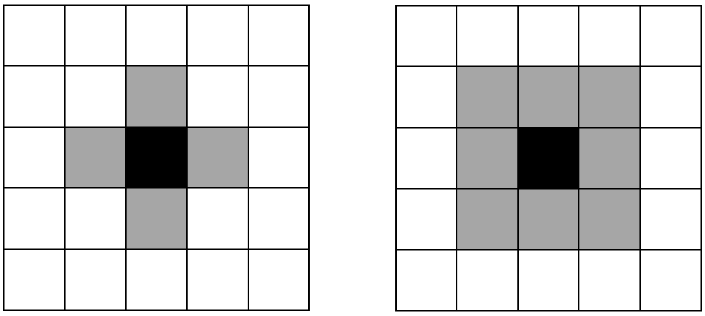

**(a)** First order von Neumann neighbourhood

**(b)** First order Moore neighbourhood

**Figure 2:** Two neighbourhoods that are commonly used in CA models.

###### 2.2.2 Cell cycle progression

Every automaton element occupied by a proliferating cell is assigned a *counter* that monitors its cell cycle progression. When a new cell is created following cell proliferation, its *counter* $\tau_{cycle_\gamma}$ is a random number drawn from a normal distribution with mean $\bar{\tau}_{cycle_\gamma}$ and standard deviation $\sigma_{cycle_\gamma}$ where $\bar{\tau}_{cycle_\gamma}$ represents the average cell cycle duration for cell population $\gamma$. After each discrete time step $\tau$, the cell cycle counter $\tau_{cycle_\gamma}$ for a given cell is reduced by an amount that depends on its local *neighbourhood* (see Fig. 3). This

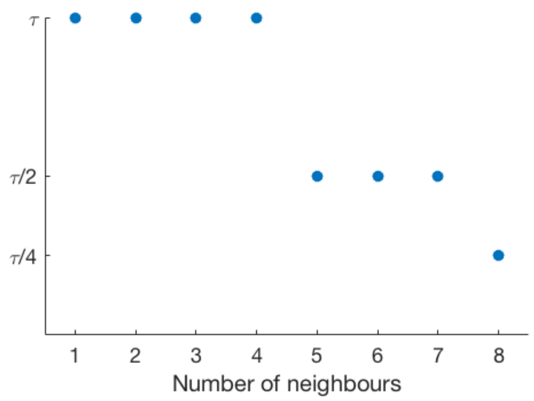


**Figure 3:** The amount by which $\tau_{cycle_\gamma}$ for a given cell is reduced after each computational step of length $\tau$ as a function of the number of the cell's neighbours.

mechanism imitates contact inhibition of proliferation, a well known feature of 2D and 3D cell aggregates. Unlike Jagiella et al. [7] who assumed that a cell cycles only if it is located within a given distance from the spheroid boundary, here a cell makes this decision based on local information only.

###### 2.2.3 Cell division

When $\tau_{cycle_\gamma} \leq 0$ for a given cell, it divides to produce two identical cells. One cell occupies the same position as its parent and the other cell is placed in an adjacent automaton element. If more than one automaton element adjacent to the dividing cell is empty then the division process is complete (if a dividing cell has more than one free neighbour then the neighbour with the maximum number of neighbours is chosen to maintain cell-cell adhesion [8]). Alternatively, if a dividing cell does not have an empty adjacent automaton element, then we find the shortest chain of cells that connects the dividing cell to the spheroid's boundary. We shift the cells in the chain along this path in order to create space for the new daughter cell (if multiple shortest paths exist we choose one at random). This algorithm mimics the way in which growing cells exert mechanical stress on their neighbours to generate spheroid expansion. Fig. 4 illustrates how the chain of cells 1, 2 and 3 is shifted towards the spheroid's boundary to create space for the new daughter cell D.

###### 2.2.4 Cell death

A cell becomes necrotic if the oxygen concentration at its location falls below a threshold value $c_{\mathcal{N}_\gamma}$. Necrotic cells are lysed at the rate $p_{lys_\gamma}$ ($\mathrm{h}^{-1}$). When a necrotic cell is lysed it is removed from the computational grid and a chain of cells is shifted from the spheroid's boundary towards the location of the removed cell

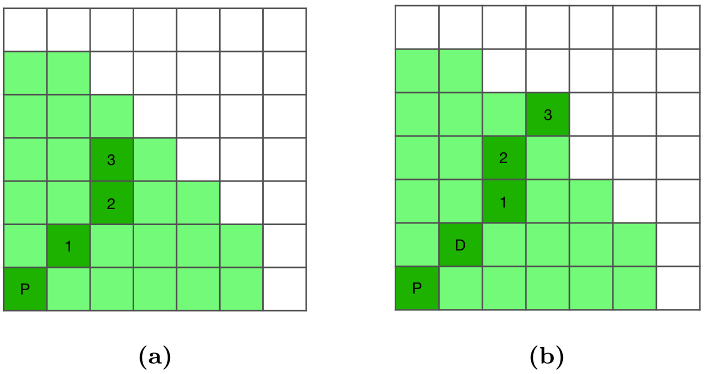


**Figure 4:** Chain shifting following a cell division event. State of the grid (a) before and (b) after division. The dividing cell P pushes the chain of cells 1, 2 and 3 toward the spheroid's boundary to create space for the new daughter cell D. Key: proliferating cells (green); empty cells (white); chain of shifted cells (dark green).

(see Fig. 5). In order to preserve spheroidicity we select the cell on the boundary that is located furthest away from the spheroid center (or choose randomly if multiple cells lie at the same distance from the center). The resulting cell rearrangement occurs within one computational time step and ensures that the spheroid remains compact.

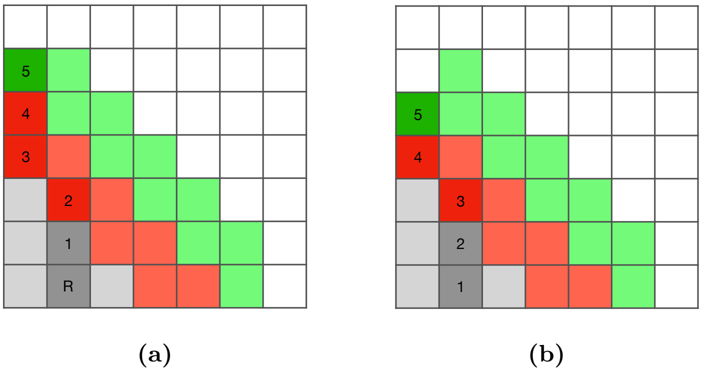


**Figure 5:** Chain shifting following lysis. State of the grid (a) before and (b) after cell removal. A chain of cells 1, 2, 3, 4 and 5 are shifted following lysis of cell R. Key: proliferating cells (green); quiescent cells (red); necrotic cells (grey); empty cells (white).

#### 2.3 Algorithm summary

We now outline the computer algorithm used to implement the hybrid CA model.

1. *Initialisation* ($t = 0$). We assign values to all model parameters (see Table 2), initialise the oxygen distribution on the spatial grid, and place a cluster of proliferating control and resistant cells at the

centre of the computational grid (all other grid sites are assumed to contain culture medium), assign cell cycle duration times to the cells and set $t = \tau$ where $\tau$ is the length of our computational step.

2. *Nutrient consumption.* We update the oxygen concentration by solving Eqs. (8)–(11).

3. *Update cell states.* We update the states of all cells based on the updated oxygen distribution (see Sec. 2.2.1).

4. *Update cell cycle times.* The cell cycle duration *counter* of each proliferating cell is decreased by an amount that depends on the number of cells in its first order Moore neighbourhood (see Fig. 3).

5. *Check for cell division.* If the updated cell cycle *counter* of a proliferating cell satisfies $\tau_{cycle} \leq 0$ then the cell divides. Cell division is performed asynchronously (i.e., we randomly loop over all cells marked for division).

6. *Check for lysis of necrotic cells.* Necrotic cells are removed from the grid at rate $p_{lys}$. Cell lysis is performed asynchronously.

7. *Update time.* Time $t$ is increased by a computational step $\tau$. If $t < T$, where $T$ is the total simulation time, then steps (2)–(7) are repeated; if $t \geq T$, then the simulation ends.

Our hybrid CA model was implemented in MATLAB. A flowchart summarising the algorithm is presented in Fig. 6.

#### 2.4 Hybrid CA model parameter estimates

##### 2.4.1 Oxygen concentration

The background oxygen concentration was set to $c_\infty = 2.8 \times 10^{-7}$ mol cm$^{-3}$ [4] and the oxygen diffusion constant within a growing spheroid was set to $D = 1.8 \times 10^{-5}$ cm$^2$ s$^{-1}$ [6]. Since the diffusion of $O_2$ in culture medium is likely to be higher than within the spheroid [7], we assumed that the concentration of $O_2$ within the medium was replenished on a faster timescale than within the spheroid. In practice this means that $O_2$ levels in the culture medium were held constant at the background $O_2$ concentration $c_\infty$. Proliferating cells were assumed to consume oxygen with rate $\kappa_{\mathcal{P}_\gamma}$. Quiescent cells also consume oxygen but at a lower rate $\kappa_{\mathcal{Q}_\gamma} = 0.5 \times \kappa_{\mathcal{P}_\gamma}$.

Measurements of oxygen consumption by the PC3 Ctrl and PC3 Res cells cultured in monolayers revealed that the Ctrl cells consume more oxygen than Res cells by a factor of 1.4 (see main text). As

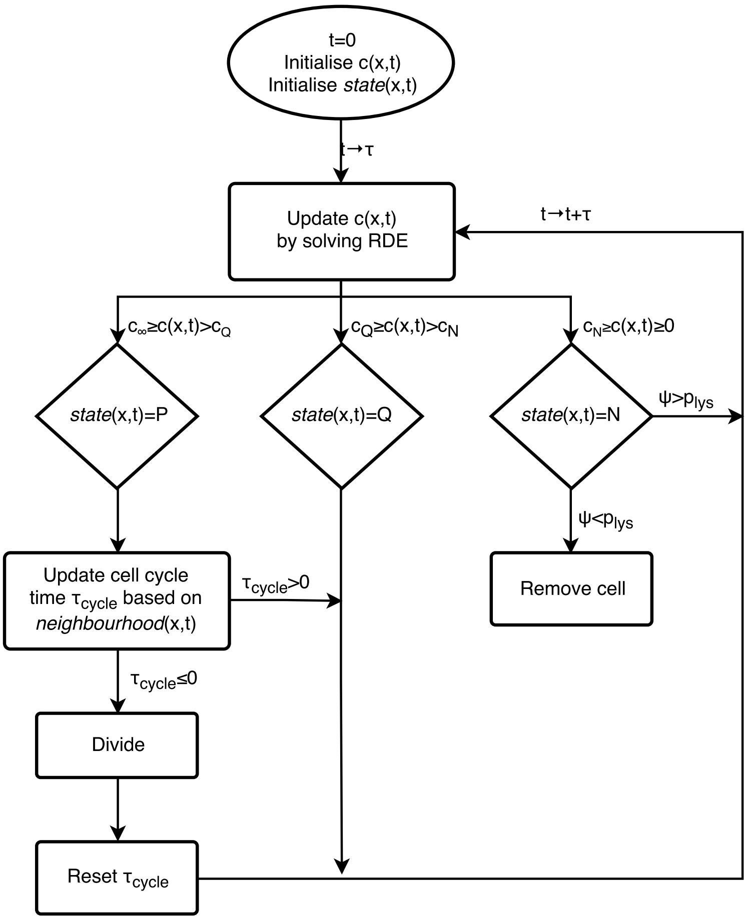


**Figure 6:** Flowchart summarising the algorithm used to implement the CA model.

shown previously, cells in tumour spheroids are known to consume less oxygen than in monolayers [4]. In the absence of estimates of the $O_2$ consumption rates in the PC3 spheroids, we treated $\kappa_{\mathcal{P}_\gamma}$ as free parameters and explored their influence on spheroid growth. Estimates of the values of $c_{\mathcal{Q}_\gamma}$ and $c_{\mathcal{N}_\gamma}$ were also difficult to obtain and so they were also treated as free parameters (see Sec. 2.4.3).

###### 2.4.2 Cell cycle duration

We estimated cell cycle times for the PC3 Ctrl and PC3 Res cell populations by seeding a small number ($N_0 = N(t=0) = 10^4$) of cells in a Petri dish, allowing them to expand for 48 hours and counting $N(t=48)$, the number of viable cells at $t = 48$ hours. By assuming that the cells grew exponentially during this time period with growth rate $k$ (so that $N(t) = N_0 e^{kt}$), we estimated the growth rate $k$ ($h^{-1}$) from the data as follows,

$$k = \frac{\log\left(\frac{N_{48}}{N_0}\right)}{48}.$$

Given $k$, the doubling time $t_d$ can be calculated via $t_d = \frac{\log(2)}{k}$ hours. We used $t_d$ to estimate the cell cycle duration time. Thus the average cell cycle time of the PC3 Ctrl was estimated to be $\bar{\tau}_{cycle_{\mathcal{C}}} = 18.3$ h with standard deviation $\sigma_{cycle_{\mathcal{C}}} = 1.4$ h. The PC3 Res cells divided more rapidly, with mean $\bar{\tau}_{cycle_{\mathcal{R}}} = 16.9$ h and standard deviation $\sigma_{cycle_{\mathcal{R}}} = 0.9$ h. These estimates were based on six independent measurements.

###### 2.4.3 Parameter inference

We used a two step process to estimate the values of parameters $c_{\mathcal{Q}_\gamma}$, $c_{\mathcal{N}_\gamma}$, $\kappa_{\mathcal{P}_\gamma}$ and $p_{lys_\gamma}$ for the PC3 Ctrl and PC3 Res cells. First, we used our hybrid CA model to simulate the growth of homogeneous spheroids for a range of values of the model parameters. We then compared the simulation results to growth curves from the PC3 Ctrl and PC3 Res homogeneous spheroids and experimental images of their spatial distributions. In this way, we identified those values of $c_{\mathcal{Q}_\gamma}$, $c_{\mathcal{N}_\gamma}$, $\kappa_{\mathcal{P}_\gamma}$ and $p_{lys_\gamma}$ that best fit the data. We then fixed these parameter values in the hybrid CA model when simulating the growth of heterogeneous spheroids.

In more detail, to identify the values of $c_{\mathcal{Q}_\gamma}$, $c_{\mathcal{N}_\gamma}$, $\kappa_{\mathcal{P}_\gamma}$ and $p_{lys_\gamma}$ that best fit the growth curves for a given cell population $\gamma \in \{$PC3 Ctrl, PC3 Res$\}$ we ran the CA model for a single population $\gamma$ for the following ranges of parameters: $c_{\mathcal{Q}_\gamma} \in (0, 2.8 \times 10^{-7})$, $c_{\mathcal{N}_\gamma} \in (0, c_{\mathcal{Q}_\gamma})$, $\kappa_{\mathcal{P}_\gamma} \in (1.0 \times 10^{-9}, 1.0 \times 10^{-8})$ and $p_{lys_\gamma} \in (0, 0.5)$. We compared the *in silico* growth curves to the experimental PC3 Ctrl and PC3 Res growth curves by calculating the sum of squared residuals between them and shortlisted ten sets of parameters that resulted in *in silico* growth curves with the lowest sum of squares for each population. We then compared the spatial distributions for the shortlisted *in silico* spheroids to the available images of homogeneous PC3

**Table 2:** Summary of the parameters used in the CA model together with estimates of their values.

| **Parameter** | **Description** | **Value** | **Units** | **Ref.** |
|---|---|---|---|---|
| $l_{PC3}$ | Cell size | 0.0018 | cm | [2] |
| $L$ | Domain length | 0.36 | cm | Estimated |
| $\bar{\tau}_{cycle_C}(\sigma_{cycle_C})$ | Mean (standard deviation) cell cycle time (PC3 Ctrl) | 18.3 (1.4) | h | Estimated |
| $\bar{\tau}_{cycle_R}(\sigma_{cycle_R})$ | Mean (standard deviation) cell cycle time (PC3 Res) | 16.9 (0.9) | h | Estimated |
| $c_\infty$ | Background $O_2$ concentration | $2.8 \times 10^{-7}$ | mol cm$^{-3}$ | [4] |
| $D$ | $O_2$ diffusion constant | $1.8 \times 10^{-5}$ | cm$^2$ s$^{-1}$ | [6] |
| $c_{Q_C}$ | $O_2$ concentration threshold for Ctrl proliferating cells | $1.82 \times 10^{-7}$ | mol cm$^{-3}$ | Estimated |
| $c_{N_C}$ | $O_2$ concentration threshold for Ctrl quiescent cells | $1.68 \times 10^{-7}$ | mol cm$^{-3}$ | Estimated |
| $c_{Q_R}$ | $O_2$ concentration threshold for Res proliferating cells | $2.24 \times 10^{-7}$ | mol cm$^{-3}$ | Estimated |
| $c_{N_R}$ | $O_2$ concentration threshold for Res quiescent cells | $2.1 \times 10^{-7}$ | mol cm$^{-3}$ | Estimated |
| $\kappa_{P_C}$ | $O_2$ consumption rate of proliferating cells | $1.0 \times 10^{-8}$ | mol cm$^{-3}$ s$^{-1}$ | Estimated |
| $\kappa_{P_R}$ | $O_2$ consumption rate of proliferating cells | $3.3 \times 10^{-9}$ | mol cm$^{-3}$ s$^{-1}$ | Estimated |
| $\kappa_{Q_\gamma}$ | $O_2$ consumption rate of quiescent cells | $0.5\kappa_{P_\gamma}$ | mol cm$^{-3}$ s$^{-1}$ | Estimated |
| $p_{lys_C}$ | Rate of lysis of Ctrl cells | 0.015 | h$^{-1}$ | Estimated |
| $p_{lys_R}$ | Rate of lysis of Res cells | 0.015 | h$^{-1}$ | Estimated |

control and resistant spheroids and selected those sets of parameter values that best matched the images for both populations by visual inspection. The selected parameter sets for the control and resistant populations were subsequently used in the simulations of heterogeneous spheroids (see Table 2).

Fig. 7 shows typical results from simulations of our hybrid CA model. Fig. 7a shows how the spatial distribution of a homogeneous spheroid changes over time for a single realisation of the model whereas Fig. 7b represents the average cumulative volumes of the necrotic, quiescent and proliferating regions from 100 realisations of the model.

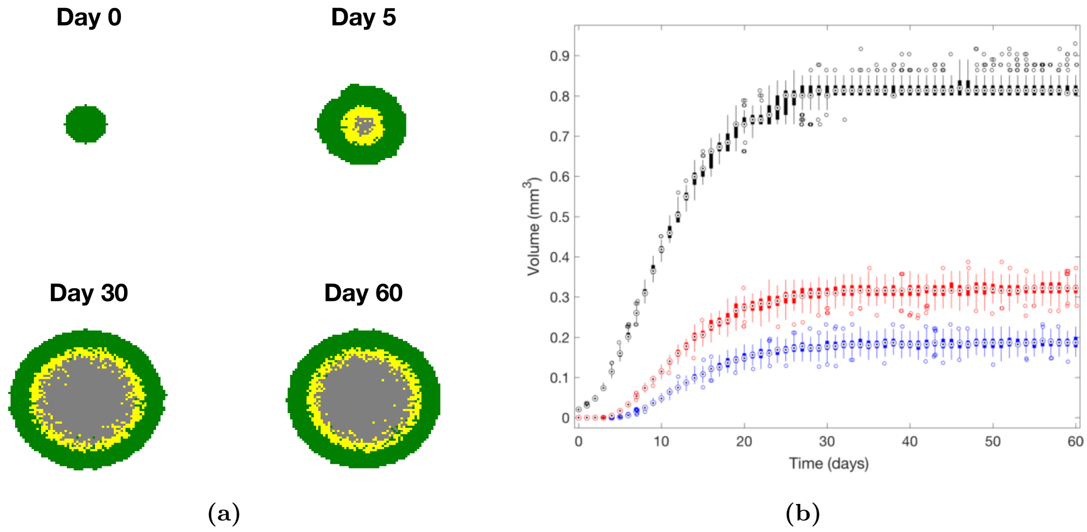


**Figure 7:** Typical results from simulations of the 2D CA model showing (a) spatial distributions of spheroids on different days from a single realisation of the model and (b) growth curves obtained from 100 realisations. In (a), the proliferating cells are green, quiescent cells are yellow and necrotic cells are grey. Plot (b) shows the total volumes of spheroids (black), the volumes of necrotic regions (blue) and the combined volumes of the necrotic and quiescent regions (red). The circled dots represent the median values, the boxes represent the interquartile ranges and the whiskers extending from the boxes mark the min. and max. values. The empty circles represent outliers. Nondimensionalised parameter values used in the simulations: $\bar{\tau}_{cycle} = 18.3$, $\kappa_{\mathcal{P}} = 150$, $p_{lys} = 0.015$, $c_{\mathcal{Q}} = 0.8$ and $c_{\mathcal{N}} = 0.775$.
